# Phase separation and molecular ordering of the prion-like domain of the thermosensory protein EARLY FLOWERING 3

**DOI:** 10.1101/2023.03.12.532276

**Authors:** Stephanie Hutin, Janet R. Kumita, Vivien I. Strotmann, Anika Dolata, Wai Li Ling, Nessim Louafi, Anton Popov, Pierre-Emmanuel Milhiet, Martin Blackledge, Max H. Nanao, Philip A. Wigge, Yvonne Stahl, Luca Costa, Mark D. Tully, Chloe Zubieta

**Affiliations:** Laboratoire Physiologie Cellulaire et Végétale, Univ. Grenoble Alpes, CNRS, CEA, INRAE, IRIG-DBSCI-LPCV, 17 Avenue des Martyrs, 38054, Grenoble, France; Dept. of Pharmacology, University of Cambridge, Tennis Court Road, Cambridge, UK, CB2 1PD; Institute for Developmental Genetics, Heinrich-Heine University, Universitätsstraße 1, D-40225 Düsseldorf, Germany; Univ. Grenoble Alpes, CEA, CNRS, IBS, F-38000 Grenoble, France; Centre de Biologie Structurale (CBS), Univ Montpellier, CNRS, INSERM, 29 Rue des Navacelles, 34090 Montpellier, France; European Synchrotron Radiation Facility, Structural Biology Group, 71 Avenue des Martyrs, 38000 Grenoble, France; Leibniz-Institut für Gemüse- und Zierpflanzenbau, Großbeeren, Germany and Institute of Biochemistry and Biology, University of Potsdam, Potsdam, Germany; Cluster of Excellence on Plant Sciences, Heinrich-Heine University, Universitätsstraße 1, D-40225 Düsseldorf, Germany

**Keywords:** Liquid-liquid phase separation, hydrogel, small angle X-ray scattering, atomic force microscopy, fluorescence microscopy, EARLY FLOWERING 3, thermosensing, Arabidopsis thaliana

## Abstract

Liquid-liquid phase separation (LLPS) is an important mechanism enabling the dynamic compartmentalisation of macromolecules, including complex polymers such as proteins and nucleic acids, and occurs as a function of the physicochemical environment. In the model plant, *Arabidopsis thaliana*, LLPS by the protein EARLY FLOWERING3 (ELF3) occurs in a temperature sensitive manner and controls thermoresponsive growth. ELF3 contains a largely unstructured prion-like domain (PrLD) that acts as a driver of LLPS in vivo and in vitro. The PrLD contains a poly-glutamine (polyQ) tract, whose length varies across natural Arabidopsis accessions. Here, we use a combination of biochemical, biophysical and structural techniques to investigate the dilute and condensed phases of the ELF3 PrLD with varying polyQ lengths. We demonstrate that the dilute phase of the ELF3 PrLD forms a monodisperse higher order oligomer that does not depend on the presence of the polyQ sequence. This species undergoes LLPS in a pH and temperature-sensitive manner and the polyQ region of the protein tunes the initial stages of phase separation. The liquid phase rapidly undergoes aging and forms a hydrogel as shown by fluorescence and atomic force microscopies. Furthermore, we demonstrate that the hydrogel assumes a semi-ordered structure as determined by small angle X-ray scattering, electron microscopy and X-ray diffraction. These experiments demonstrate a rich structural landscape for a PrLD protein and provide a framework to describe the structural and biophysical properties of biomolecular condensates.

## Introduction

Compartmentalisation into biomolecular condensates, or membraneless organelles, helps to regulate the biochemistry of the cell by dynamically concentrating and sequestering different components including proteins such as transcription factors, RNA-binding proteins and co-factors and nucleic acids ^1–9^. Proteins with intrinsically disordered regions (IDRs) and low complexity prion-like domains (PrLD) often act as drivers of LLPS, separating into a highly concentrated protein-rich phase and a dilute phase under specific conditions^10–12^. While simple polymers have been successfully studied experimentally and modelled using theoretical methods such as course-grained (CG) simulation and atomistic models, quantifying and predicting the behaviour of PrLD proteins as a function of physicochemical variables is challenging due to the complexity in the amino acid sequence of proteins^13–19^. The interactions that contribute to the metastable condensed phase include many transient, short-range interactions including pi-pi, cation-pi, dipole, electrostatic and hydrophobic interactions, all of which may be present in a given polypeptide. These weak and low-specificity contacts are often present in disordered proteins and will occur intramolecularly in the dilute phase and both intra- and intermolecularly in the condensed phase. The formation and dynamics of phase separation mediated by PrLD proteins is often highly sensitive to pH, ionic strength and temperature and will vary as a function of the properties of the amino acids (i.e. polar, charged, hydrophobic, aromatic) in the PrLD sequence^20–23^. While this dynamic response of PrLD proteins is technically challenging to study, it may play a critical physiological function, allowing PrLD proteins to act as sensors of the in vivo cellular environment and to alter physiological responses accordingly. For example, the poly(A)-binding protein (Pab1) in yeast and the recently characterised circadian clock protein EARLY FLOWERING3 (ELF3) in Arabidopsis have been shown to act as direct in vivo temperature sensors and to alter developmental responses^6,24^. Thus, an understanding of the molecular basis of environmental sensing by PrLD proteins is a pre-requisite to engineering these properties for the creation of tailored responses to stresses such as temperature changes in the cell.

ELF3 displays a well-characterised ability to undergo LLPS in vitro and in vivo and exists with natural sequence variation within the PrLD associated with specific phenotypes, suggesting a physiological role for LLPS ^6,26^. ELF3 is a largely disordered protein with a C-terminal PrLD. The PrLD contains a poly-glutamine repeat (polyQ) that exhibits different lengths from 7 to 29 glutamines across 181 natural Arabidopsis accessions^25,26^. Previous experiments in Arabidopsis plants grown at 17 °C, 22 °C and 27 °C demonstrated that the length of the polyQ has a mild but statistically significant effect on hypocotyl elongation, a commonly used measure of thermoresponsive growth^6^. While ELF3 PrLD with 7 glutamines (Q7), from the lab strain Columbia-0, has been shown to phase separate in vitro in a temperature-dependent manner, the effects of varying polyQ length on the dynamics of phase separation is not known^6^. In order to better understand the biophysical basis of ELF3 condensation, we verified that ELF3 Q0, Q7 and Q20 could form puncta in vivo and performed biochemical, structural and biomechanical studies of the corresponding PrLD regions, demonstrating that LLPS of ELF3 PrLD with varying polyQ lengths is sensitive to temperature and pH. We further show that the dilute phase, unlike a canonical intrinsically disordered protein, exists as a monodisperse higher order oligomer and that upon liquid-liquid phase separation forms a new species with distinct microenvironments and different biomechanical properties. This species is able to further undergo aging into an ordered hydrogel. ELF3 PrLD exhibits a distinct structural landscape, with the dynamics and biomechanical properties of the condensed liquid and gel phases modulated by the length of the polyQ region.

## Results

### In vivo puncta formation of ELF3 Q0, Q7 and Q20

Based on our previous studies of the full-length and PrLD of ELF3, the protein undergoes LLPS in vivo and in vitro with the PrLD required and sufficient for phase separation^6^. To verify formation and dynamics of condensate formation in vivo, the sequence of the *ELF3* gene encoding polyQ lengths of 0 (deletion mutant), 7 (from accession Columbia-0) and 20 (from accession Sandåkra-2 (San-2)) glutamines tagged with mVenus was expressed under an inducible promoter in tobacco leaf epidermal cells. The formation of puncta was observed for all three constructs (Figure 1). Fluorescence recovery after photobleaching (FRAP) experiments were performed to investigate the dynamics of the proteins in the puncta. The puncta for all constructs exhibited partial recovery after photobleaching, with fluorescence intensities after photobleaching ranging from about 50% for Q0 and Q7 (Figure 1A and B) to 30% for Q20 (Figure 1c) of the prebleach intensity. This indicates that the puncta are comprised of a mixture of mobile and immobile species, as has been observed for many other systems^27^. In order to more robustly characterize the dynamics and structure of the condensate, detailed in vitro studies of the PrLD region of the protein required for phase separation were performed.

**Figure 1.**
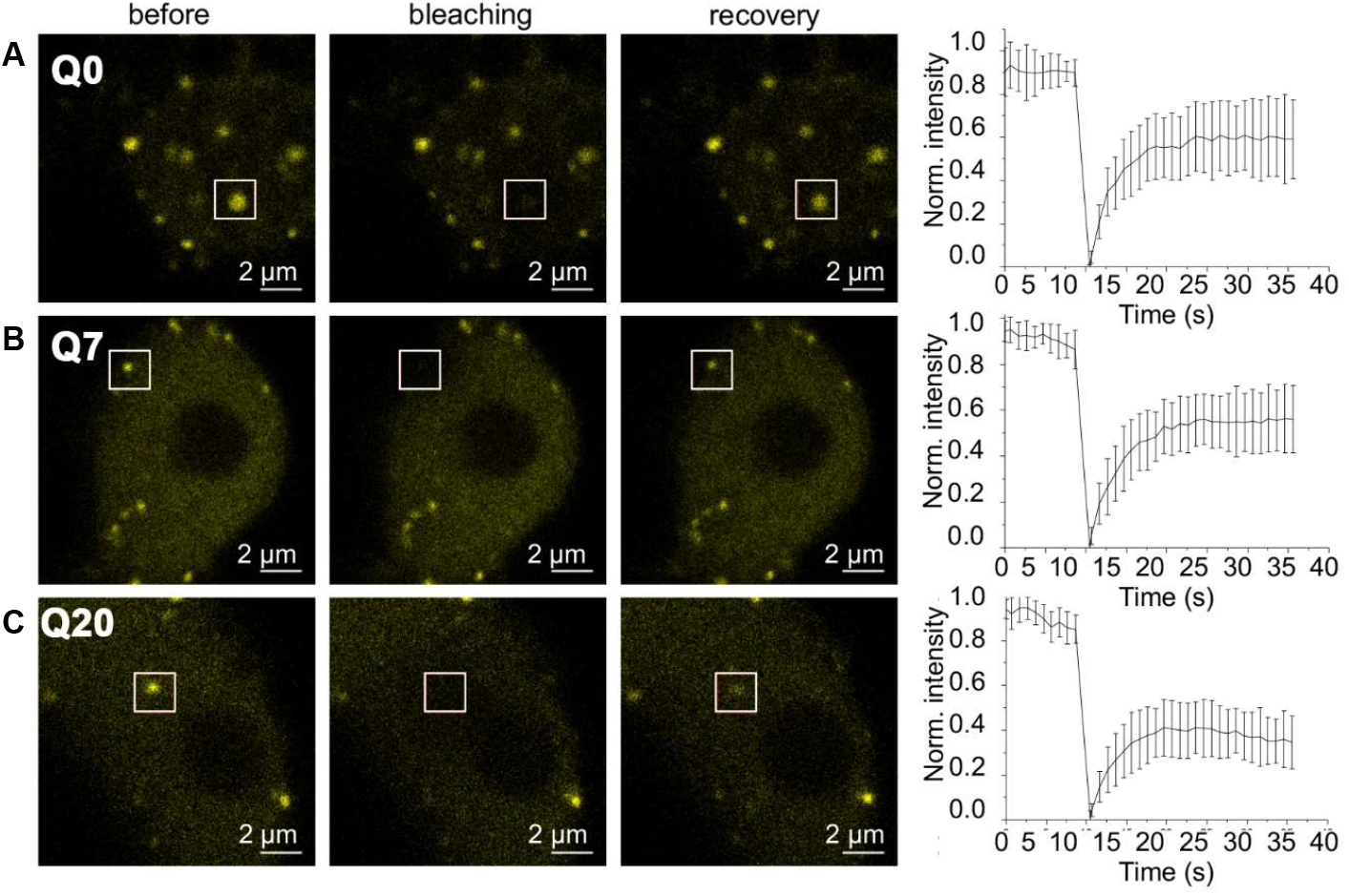
Representative fluorescence recovery after photobleaching (FRAP) images of agroinfiltrated *Nicotiana benthamiana* leaf epidermal cells transiently expressing mVenus labelled ELF3 Q0, Q7 or Q20. White boxes indicate photobleached areas. Panels at left are prior to bleaching, middle immediately after bleaching and right after recovery. Mean recovery curves for ELF3 Q0, Q7 and Q20 are shown at far right and S.D. are shown. All curves are generated from 9-10 individual puncta measurements with each FRAP experiment performed on a different cell.

### In vitro characterization of the ELF3 PrLD

Using optical imaging, turbidity assays (A440) and phase diagrams, we examined the effects of temperature, protein concentration and pH on LLPS of ELF3 PrLD with polyQ lengths of 0, 7 and 20 glutamines (Figure 2 and Supplementary Figure S1A). All PrLD constructs underwent LLPS, however temperature and pH had subtle but measurable effects on phase separation for the three constructs studied. Increasing temperature triggered LLPS for Q0, Q7 and Q20 constructs and exhibited reversibility as shown qualitatively (Figure 2A) using temperature steps. Absorbance measurements at A_440_ were used to further examine this behaviour, with Q0 and Q7 behaving similarly, exhibiting an LLPS transition temperature occurring at 31.2 ±0.4 °C and 28.1±0.3 °C, respectively (Figure 2B). In comparison, the Q20 construct started phase separation at a lower temperature, as shown by the non-zero normalised absorbance between 15 °C and 20 °C and a sloping bottom plateau. The A_440_ gradually increased for Q20 as the temperature was raised with a Tm of 33.0 ±1.5 °C (Figure 2B). The Tm values were calculated based on the first temperature ramps, as denoted by the filled blue squares in Figure 2b. The effects of temperature were largely reversible for all polyQ constructs tested, with the PrLD proteins switching between the dilute and condensed phase as monitored by A_440_ (Figure 2A and B). It should be noted that, due to the experimental set-up for the A_440_ measurements, where the highest temperature measured was 40 °C, the parameters extracted from the fitting, in particular for Q20, should be considered as estimations as no plateau was reached and above 40 °C the proteins began to irreversibly precipitate. The observed demixing of the ELF3 PrLD proteins with increasing temperature is indicative of hydrophobic and aromatic interactions driving phase separation due to the favourable enthalpic contributions to the free energy of the system under the temperature regime of interest and positive entropic effects due to counter ion and/or hydration water release, as has been observed for other LLPS systems^20,28–30^. All constructs underwent phase separation under the conditions tested and expansion of the polyQ repeat from 0 to 20 glutamines had a relatively small effect on the transition temperature onset of phase separation.

**Figure 2.**
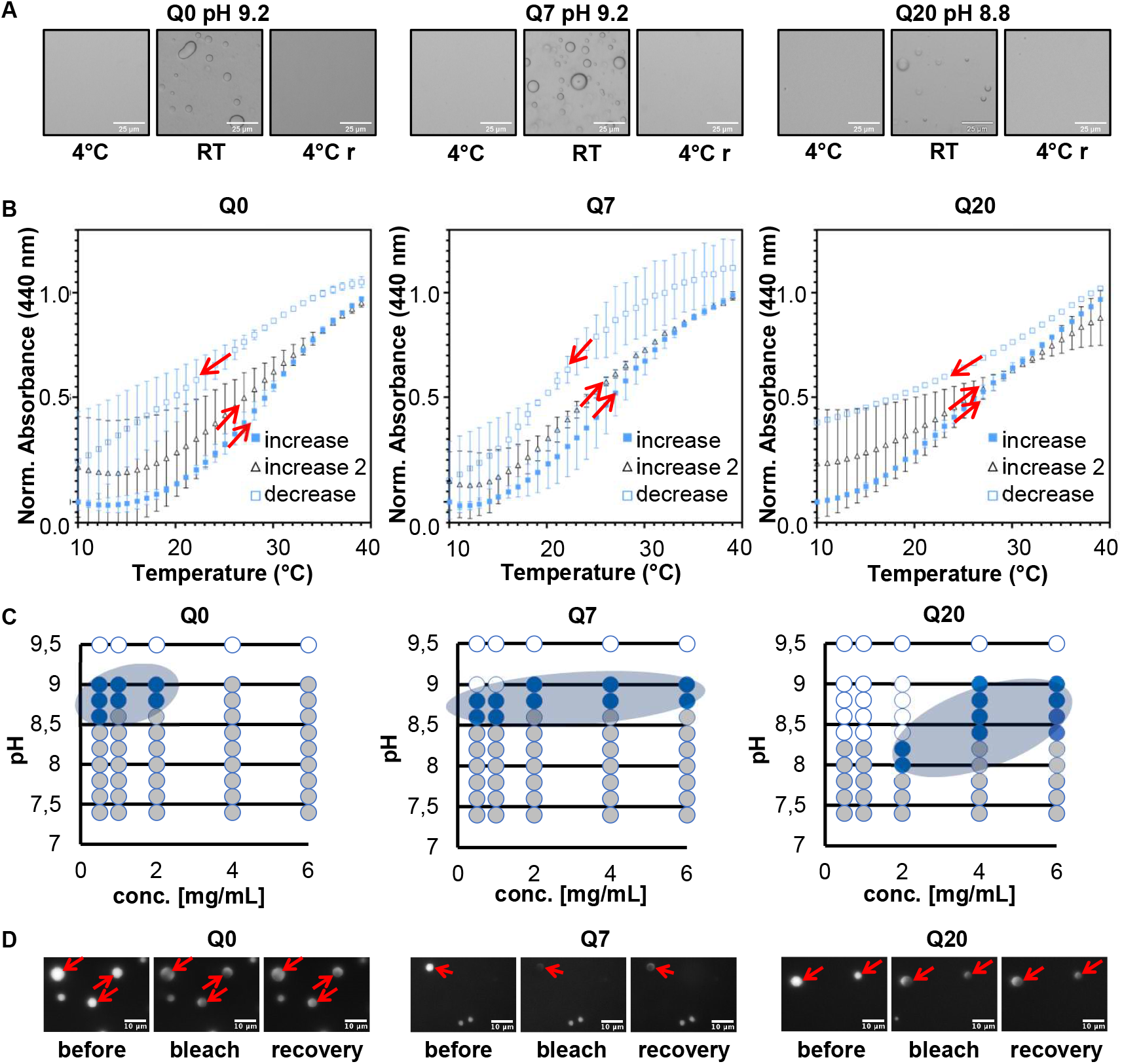
ELF3 PrLD with varying polyQ lengths undergoes phase separation. **A)** Reversible temperature induced phase separation for ELF PrLD Q0, Q7 and Q20 (4 mg/mL) visualised at 4°C (left), warmed to 22°C (center) and cooled to 4°C (right). **B)** Turbidity assays (A440) of untagged ELF3 PrLD Q0, Q7 and Q20 as a function of temperature. Protein concentration was 15μM (~0.4 mg/ml) for each construct. Temperature ramps are shown with red arrows indicating increasing or decreasing temperature. **C)** Phase diagrams for untagged ELF3 PrLD Q0, Q7 and Q20 constructs as a function of protein concentration and pH. Open circles are dilute phase, dark blue LLPS and grey precipitate/gel. LLPS conditions are shaded in light grey for clarity. Circles represent measurements at a specific pH (9.5, 9.0, 8.8, 8.6, 8.4, 8.2, 8.0, 7.8, 7.6 and 7.4) and a specific protein concentration (0.5, 1.0, 2.0, 4.0, 6.0 mg/ml) **D)** Fluorescence recovery after photobleaching (FRAP) experiments. Red arrows indicate regions that were photobleached and scale bars are shown. Recovery panels displayed are 3 minutes after photobleaching for all samples. Samples exhibited little fluorescence recovery due to formation of an immobile gel phase.

In addition to temperature variables, pH changes also affected phase separation, with the longer polyQ construct (Q20) forming spherical droplets over a wider range of protein concentrations and pH (pH 8.0-9.0) than the Q0 and Q7 constructs, which only underwent LLPS over a narrow pH range of 8.5-9.0 under the same ionic strength buffer conditions (Figure 2C). Fusing a C-terminal GFP to the PrLD constructs resulted in the formation of spherical droplets with the same overall trend observed for the unfused proteins, with the Q20 construct undergoing phase separation at a pH range of 6.5-8.5 versus the Q7 and Q0 constructs that behaved in a similar manner, with phase separation occurring between pH 7-8.5 (Supplementary Figure S1B). Extending the polyQ region results in a broader range of LLPS formation with respect to pH and temperature. This may be due to the relatively short polyQ tracts for Q0 and Q7, with longer polar glutamine stretches acting to prevent precipitation and keeping the protein in the condensed droplet phase over a wider range of pH and temperatures.

### Fluorescence recovery after photobleaching in vitro

While pH and temperature affected LLPS, we questioned whether there were inherent differences with respect to dynamics or stability of the condensed phase due to varying polyQ length. Fluorescence recovery after photobleaching (FRAP) experiments were performed on Q0, Q7 and Q20 PrLD constructs fused to GFP to investigate this possibility. All proteins formed droplets as the pH of the solution was reduced step-wise (0.2 pH units) from 9.4 to 8.4 and 40 minutes of equilibration after each pH change. These experiments revealed the formation of a low mobility condensate with negligible fluorescence recovery over minutes (Figure 2D). No clear differences were observed for the different samples, with all samples exhibiting formation of gel-like droplets. This suggests that while temperature change results in more reversible liquid-liquid phase separation, at least under fast temperature ramps, pH changes favoured gel formation after initial condensation.

### Biomechanical measurements by AFM

To further characterise the condensed phase and determine whether there were differences in the biomechanical properties of the samples, atomic force microscopy (AFM) and AFM-coupled confocal microscopy^31^ experiments were performed (Figure 3 and Supplementary Fig. S2-S3). All samples were applied to a glass coverslip and imaged in liquid for AFM topography and force curve measurements. Lower pH and longer adhesion times resulted in the formation of the gel state whereas higher pH and shorter adhesion times maintained the droplets in a more liquid-like state and droplets were considered a “highly viscous liquid” if they exhibited fusion with other droplets and at least 10% fluorescence recovery over 2 minutes. For these samples, in agreement with models describing nanoscale indentation of liquid interfaces,^32,33^ the stiffness was calculated based on a linear fit to the force vs. distance curves with between 9 and 13 measurements taken from the centre of each droplet to avoid edge artefacts, which are present when the droplet size is comparable to the size of the AFM tip height (≈ 3.5 - 7 μm). Such a force versus indentation linear regime can directly be related to the liquid-liquid interfacial tension^33^. This linear model is in good agreement with the data versus parabolic models for isotropic elastic solids. The droplets for all samples exhibited variable mechanical properties, suggesting the presence of harder and softer regions within individual droplets, with measurements ranging from 3 to 9 mN/m (Table 1 and Figure 3). This is likely due to liquid-hydrogel phases coexisting in the droplet and the more hydrogel present, the greater the deviation from linearity of the force as a function of distance. In contrast to the liquid condensed phase, the gel-state exhibited no fluorescence recovery, no fusion and a non-linear force versus indentation regime that could be fit using a Hertz model (applicable to solid samples) and the Young’s modulus calculated^33^ (see comparison between indentation cycles performed on liquid-like and hydrogel droplets in Supplementary Figure S2C, were deviation from linearity in the liquid phase curve can also be due to the presence of an out of contact Debye’s screening length region^33^). Interestingly, gel transition of the samples only partially affected overall droplet morphology since all droplets retained a quasi-spherical shape (Figure 4), however indentation cycles showed a non-linear force *vs*. distance regime as opposed to the linear force *vs*. distance pattern observed for the highly viscous liquid droplet samples. In addition, the mechanical response changed for the hydrogel samples. For all hydrogel samples, the stiffness ranged between 5-100 kPa, with ELF3 PrLD Q0 showing a consistently lower Young’s modulus than the Q7 and Q20 constructs (Table 1). It should be noted that constructs in Figure 4A exhibit higher rigidity than the values reported in Table 1 because they were imaged with a higher AFM loading rate (see Experimental Methods), required to decrease the acquisition time while providing sufficient pixels for high resolution, thus the apparent “stiffer” response at higher loading rate is due to viscoelastic behaviour. Wide-field fluorescence and high-resolution AFM imaging of the samples further revealed a heterogeneity within the droplets, suggesting the formation of discrete microenvironments (Figure 4). The microenvironments exhibited a stacked or layered structure, with varying step-like height profiles ranging from ~20 nm to 200 nm (Figure 4D and Supplementary Figure S3). These experiments represent the first examples, to our knowledge, of the use of AFM to determine the biomechanical properties of macromolecular condensates in the liquid phase, allowing us to probe the molecular organisation of molecules within individual droplets.

**Figure 3.**
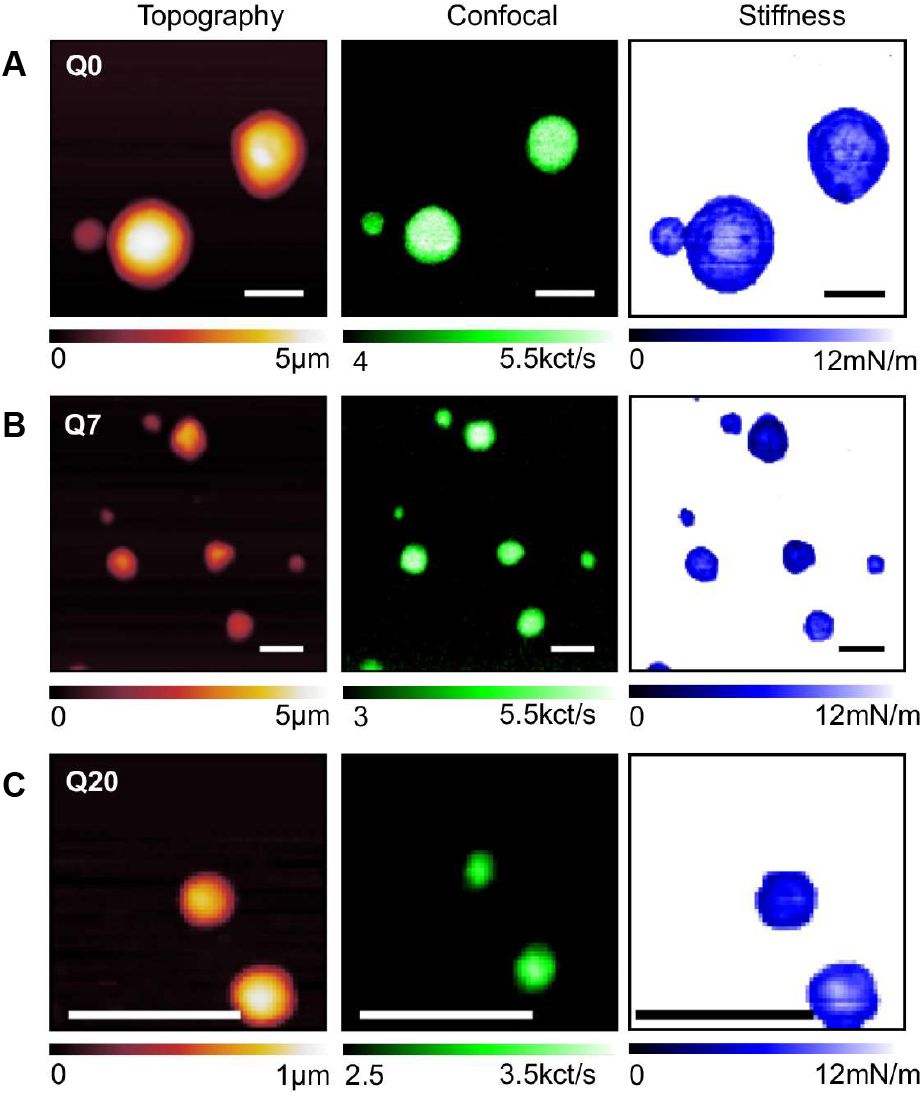
Coupled AFM and confocal fluorescence microscopy of ELF3 PrLD -GFP constructs. AFM topography mapping is shown at left, fluorescence of the same sample is shown at middle and stiffness measurements are shown at right. Alignment between the AFM and confocal microscope was within 1-2 microns for the two microscopes. Scale bar = 5 μm.

**Figure 4.**
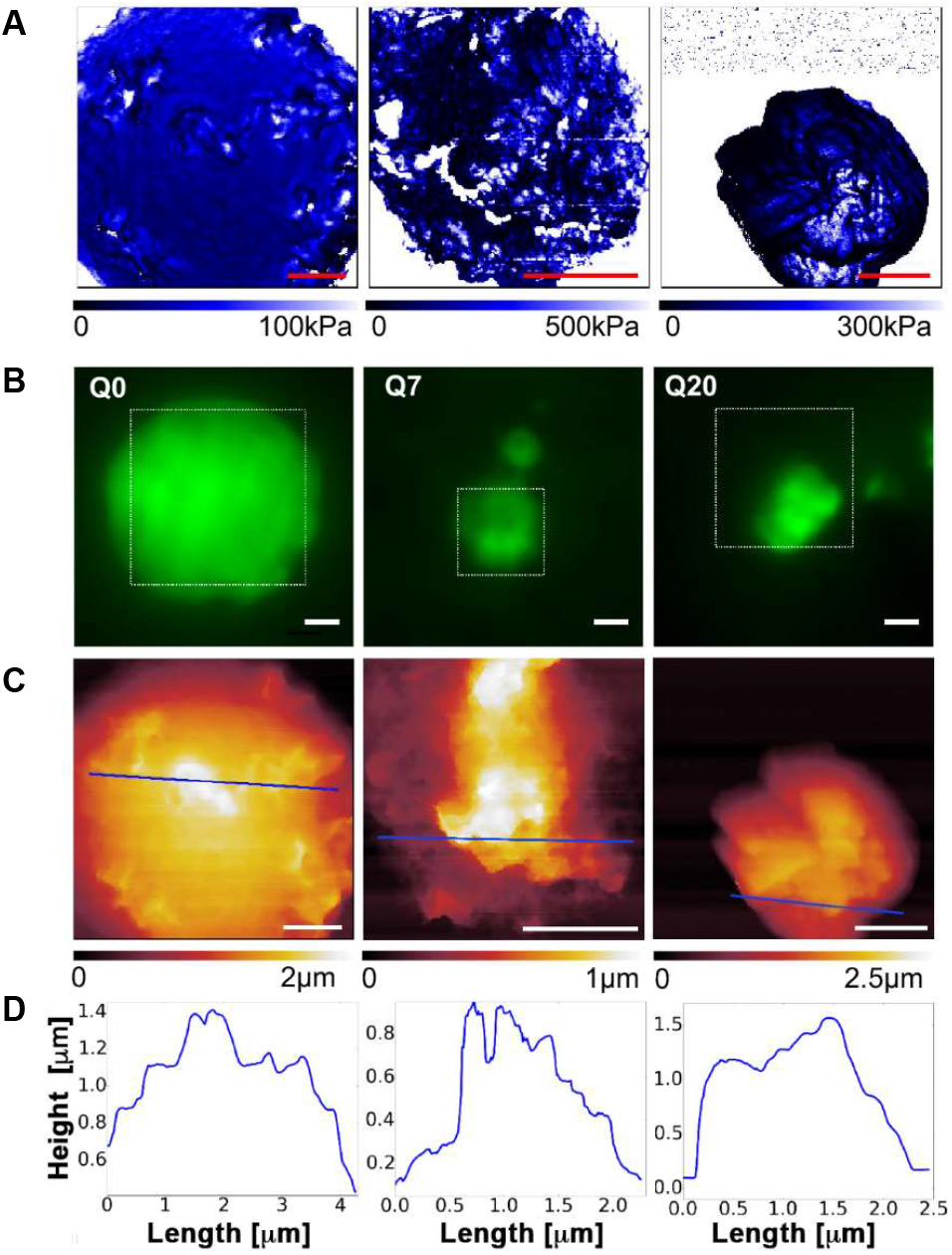
Wide-field total internal reflection fluorescence microscopy and high-resolution AFM of ELF3-GFP PrLD gel droplets. **A)** Young’s modulus measurements of the droplets showing in-homogeneous stiffness within individual droplets, scale bar = 1 μm. **B)** Fluorescence images of ELF3 PrLD Q0 (left), Q7 (middle) and Q20 (right), scalebar = 1 μm. Areas of interest are boxed and shown in close-up in (c). **C)** AFM topography of the droplets in a, scalebar = 1 μm. **D)** Height profile measurements across individual droplets (blue lines in c) showing a step pattern between flat layers.

### SAXS of dilute and condensed phases

In order to further investigate the transition from the dilute to condensed phase, condensed phase dynamics and the potential structuration and ordering of microenvironments within the condensate, we performed a series of small angle X-ray scattering (SAXS) experiments (Supplementary Table 1 and Figure 5). Scattering curves were determined for each sample (ELF3 PrLD Q0, Q7 and Q20) in the dilute phase using an online HPLC purification step in order to remove any aggregates (Figure 5A). All proteins eluted as symmetrical peaks after the void volume of the column. In the dilute phase with monodisperse samples, there should be little or no contribution to the scattering from other species such as aggregates or the condensed phase, and the scattering curve will then reflect the form factor, P(q), providing information on the size and shape of the individual molecules in solution. Based on polymer theory, the conformation the molecules adopt in the dilute phase can be predictive of their phase behaviour, providing a technically simple way to gain insight into the condensed phase^34^. Calculating the radius of gyration (Rg) based on fitting the linear portion of the Guinier region, yields values between 72-76 Å for the three samples (ELF3 PrLD Q0, Q7 and Q20) and the maximum dimension, D_max_, values of 272, 273 and 294 Å, respectively. Surprisingly, based on these measurements the estimated molecular weights, even in the dilute phase, of the molecular species corresponds to a multimeric ~28-30-mer assembly (Supplementary Table 1). This higher order oligomeric state was further confirmed by size-exclusion chromatography multiangle laser light scattering (Supplementary Figure S7). As expected, the largest Rg and D_max_ values were calculated for ELF3 PrLD Q20 due to its increased length from expansion of the polyQ region (Table 2). Furthermore, these dilute phase measurements demonstrate that for all PrLDs, the species in solution is relatively globular, as shown in the Kratky plot (Figure 5B). The compact spherical structure of a monodisperse higher order oligomer, in contrast to an extended conformation often observed for intrinsically disordered proteins, coupled with the lack of predicted secondary or tertiary structure, suggests that the presence of weak multivalent interactions may be sufficient to exclude solvent and that hydrophobic amino acid sequences lacking predicted secondary structure, partially organise the protein molecules, likely a prerequisite for phase separation by ELF3^35^. As liquid-liquid phase separation requires multivalent interactions, the dilute phase organisation may recapitulate these types of interactions intramolecularly or as smaller oligomeric units, with proteins predicted to phase separate potentially exhibiting a more compact structure versus an extended conformation from non-phase separating disordered proteins^35^.

**Figure 5.**
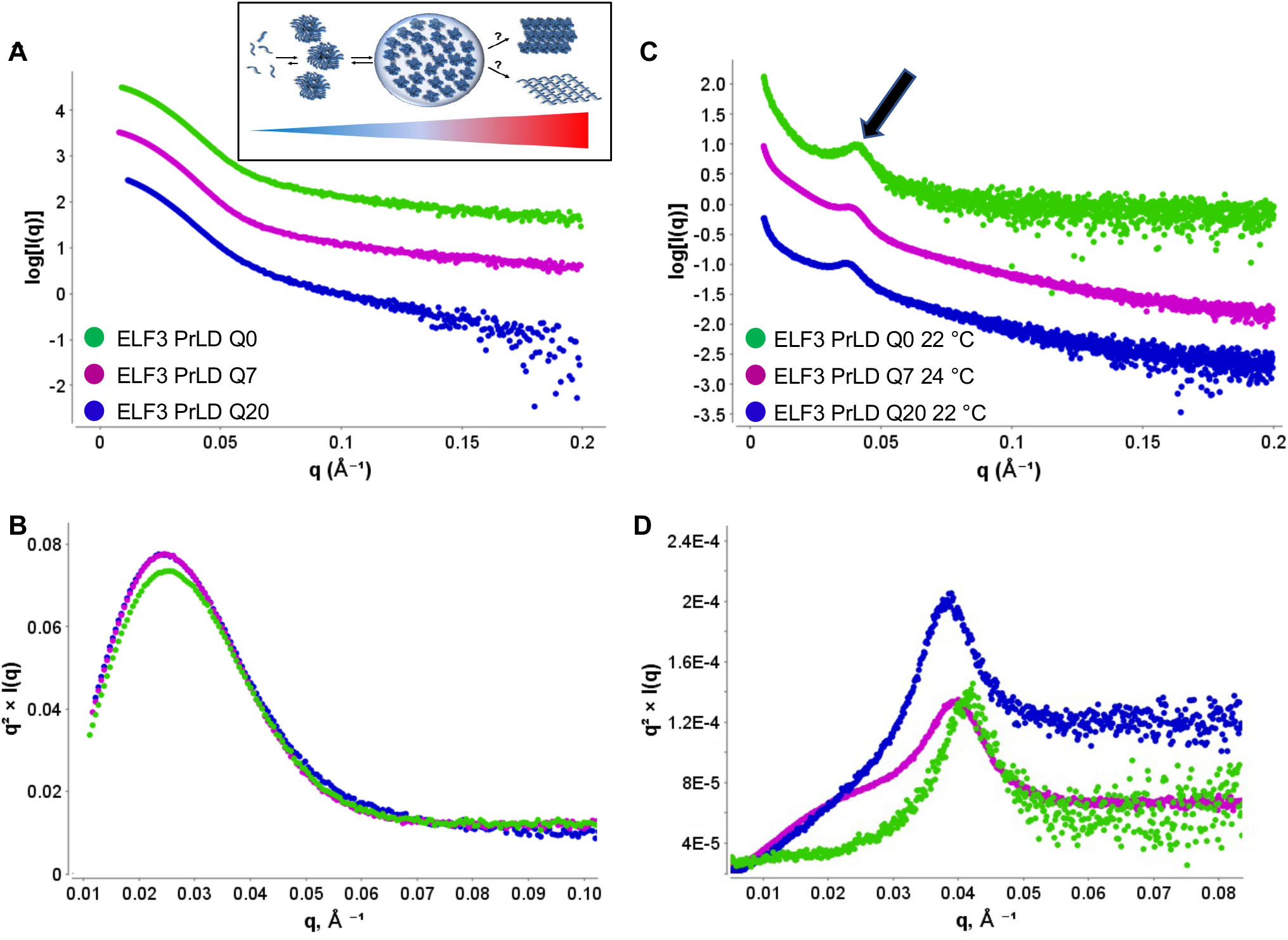
ELF3 PrLD in the dilute and condensed phase. **A)** SAXS scattering curve for ELF3 PrLD Q0 (green), Q7 (pink) and Q20 (dark blue) in the dilute phase measured with online HPLC purification. Inset shows a schematic representation of the dilute and condensed phases as a function of temperature. **B)** Corresponding Kratky plots for the curves shown in **(A)** The Gaussian shape denotes a globular species in solution. **C)** SAXS scattering curves for ELF3 PrLD Q0 (green), Q7 (pink) and Q20 (dark blue) samples in the condensed phase after aging. The structure factor peak due to long-range ordering of molecules within the condensed phase is indicated by an arrow. **D)** Corresponding Kratky plots for condensed phase scattering curves are shown and colour coded as above. The shape of the Kratky plot indicates a more extended and less globular species with the structure factor peak showing more prominently. SAXS scattering curves in **(A)** and **(C)** show a vertical offset by a factor of 10 for ELF3 PrLD Q7 and a factor of 100 for Q0 to aid in viewing.

LLPS of the samples was triggered by a temperature increase and resulted in changes to the scattering curves (Figure 5C and Supplemental Figure S5). In the liquid-liquid separated phase, the SAXS measurements demonstrate an increase in the Rg and a more extended conformation based on the Kratky plot calculations (Table 2 and Figure 5D). Changes in Rg in the condensed state have been previously observed for proteins undergoing LLPS and ascribed to nucleation events^35^. The LLPS samples are more heterogeneous than the online HPLC purified samples in the dilute phase, as reflected in the upturn of the scattering curve as I(q) approaches 0 (Figure 5C), but exhibit a clear trend in Rg, with an increase in the Rg in the condensed phase, suggesting possible elongation of the polypeptide chains or an increase in the overall size. In addition to these changes in the Rg, a structure factor peak, S(q), formed in the low q region of the scattering curves that increased in height with increasing temperature (Figure 5C and Supplemental Figure S5). This S(q) (structure factor) peak is indicative of loose lamellae within the condensed phase, and a calculated spacing (d_hk1_) of approximately 155, 163 and 167 Å for ELF3 PrLD Q0, Q7 and Q20, respectively, similar to the calculated layer spacing observed in AFM experiments. It should be noted that the layered arrangement is distinct from the characteristic fibril formation observed for amyloids,^36^ but a less compact lamellar phase characteristically observed for lipid membranes and polymer lamellae ^37,38^. To further investigate the temperature effects to the condensed phase, temperature ramps ELF3 PrLD Q7 and Q20 exhibited an increase in the structure factor peak as the temperature was increased, whereas ELF3 PrLD Q0 exhibited little change, suggesting it had passed through the liquid phase and had undergone aging to the more stable hydrogel, losing the capacity to easily transition between the condensed and dilute phases (Supplemental Figure S5).

### Structural characterization of the gel phase

The ordered species in the condensate were further characterized using stable gel samples, obtained by aging of the condensed liquid phase at pH 7.7-8 for ~1 hr at 4 °C for Q0, Q7 and Q20. These samples no longer exhibited reversibility between dilute and condensed phases with temperature or pH change and coalesced into large non-spherical amorphous species. Gel samples were characterised using transmission electron microscopy (TEM) and X-ray diffraction. Negative stain TEM confirmed regions with a stacked structure in the gel phase for all polyQ constructs, with a stack spacing estimated at ~40-50 Å (Figure 6A). This stack spacing is smaller than observed for AFM and SAXS experiments and may be due to the longer aging of the sample and/or dehydration effects of sample preparation, however a highly ordered organisation of molecules is observed. The TEM experiments exhibit inhomogeneity in the samples with certain regions exhibiting well-ordered molecular stacks and other regions showing no clear organisation.

**Figure 6.**
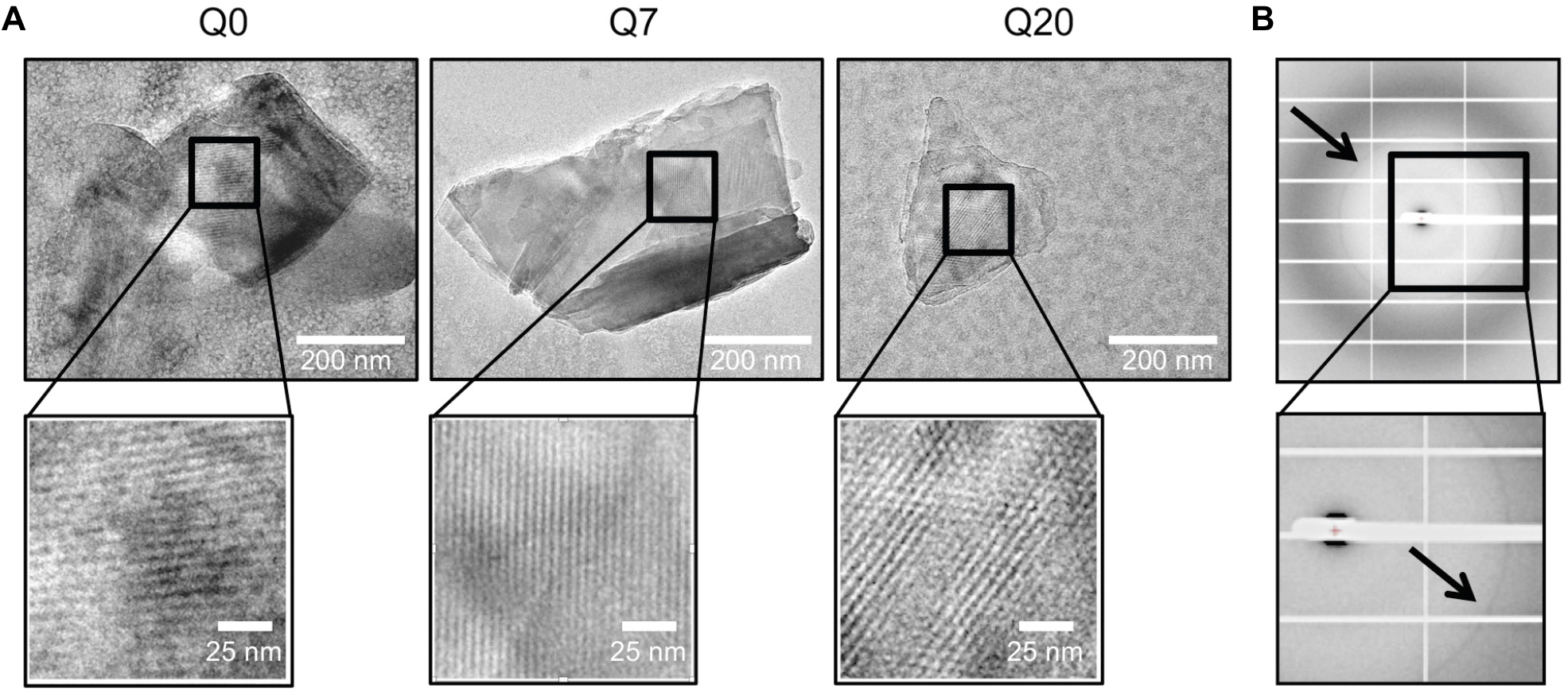
ELF3 PrLD exhibits stacked molecular organisation in the condensed phase. **A)** Negative stain TEM images of ELF3 PrLD Q0, Q7 and Q20 showing a layered structure in the gel condensed phase. Layered structure can be clearly observed in magnified images of the boxed regions. **B)** X-ray diffraction image for ELF3 PrLD Q20 showing a diffuse powder ring at 4-5 Å. A close-up is shown with the diffraction ring indicated by the black arrow.

X-ray diffraction experiments for the ELF3 PrLD Q20 construct exhibited a weak powder diffraction ring at 4-5 Å (Figure 6b), similar to observations for hydrogel forming FUS constructs, denoting a long-range molecular ordering with a powder diffraction ring arising from the distance between sheets of polypeptide chains^39^. The ELF3 PrLD Q0 and Q7 constructs showed very diffuse scattering in the 4-5 Å range (Supplemental Figure S6), due to either diffraction measurements from an amorphous region exhibiting less molecular stacking or indicating an overall less well-ordered hydrogel for the ELF3 PrLD Q0 and Q7 constructs.

## Discussion

Proteins that drive LLPS are under active study not only due to their key biological role in cellular compartmentalisation, but also for diverse applications in biomaterials and drug-delivery^20,40–44^. Characterisation of the dilute and condensed phases and the effects of exogenous variables including temperature and pH is experimentally challenging due to the complexity of the protein polymer and the intra- and intermolecular interactions at play during the nucleation and condensation process. Here, we demonstrate the utility of complementary biophysical and structural techniques to determine the dynamics and structure of different phases of a PrLD protein as a function of pH and temperature. Phase diagrams, A_440_ assays and SAXS measurements demonstrate that the dilute phase is a higher order oligomer and formation of the condensed phase is triggered by pH changes and an increase in temperature. While phase separation of the ELF3 PrLD does not depend on the polyQ regions, variation in polyQ length, which occurs in natural Arabidopsis accessions, alters LLPS formation with respect to temperature and pH. Upon aging, an ordered low mobility gel phase quickly forms in vitro and can also be observed in vivo, as puncta do not exhibit full fluorescence recovery. AFM, SAXS, EM and X-ray diffraction of the gel phase are all consistent with a stacked/layered structure in the biomolecular condensate, with extensive ordering occurring in the aged hydrogel. The physiological function of these different phases remains an open question requiring further investigation. However, with these detailed structural and biophysical studies the oligomeric state and the demixing properties of the protein can measured and quantified, providing a foundation for altering these variables and determining their physiological function.

The varying conformations and dynamics of the ELF3 PrLD polypeptide- from a surprisingly large and monodisperse oligomeric species in the dilute phase, to a solvent excluded liquid condensed phase and finally to a highly ordered hydrogel - demonstrate how the intrinsically disordered polypeptide is able to access different regions of structure space as a function of the physicochemical environment. The low-affinity and transient interactions of the ELF3 PrLD, consisting of both intra- and intermolecular interactions, may facilitate the priming of the molecules for phase separation and condensation via the formation of a large oligomeric species in the dilute phase^45,46^. While the ELF3 PrLD has little predicted secondary structure, it is still able to form a defined and homogeneous ~28-30 oligomeric assembly in solution. Based on the SAXS studies, this species is compact and globular, likely serving as an intermediate for phase separation which would occur via the fusion of oligomeric spheres. Whether this state is specific to the ELF3 PrLD or more common to PrLD phase separating proteins is unknown and will require further study.

Examination of the primary amino acid sequence of ELF3 PrLD demonstrates a number of proline clusters and aromatic residues that may act as “stickers” for LLPS and/or oligomerisation, although the specific amino acids driving this remain to be determined. In other systems that undergo LLPS and fibril or hydrogel formation, various motifs important for steric zippers, such as those in amyloid fibrils, and kinked beta sheets have been identified. Studies of LLPS in the protein FUS, for example, reveal the presence of specific peptide motifs that act as drivers for phase separation and hydrogel formation. These low-complexity, aromatic-rich, kinked segments (LARKs), have been shown to self-assemble into stacked structures and protofilaments, with short LARK hexapeptides able to form kinked beta sheets that are less stable than beta amyloid fibrils but still exhibit long-range ordering^39,47^. The ELF3 PrLD does not possess canonical LARK peptide sequences and increasing temperature generally disrupts LARK-type peptide interactions, in contrast to increasing temperature triggering LLPS for ELF3 PrLD, suggesting other mechanisms are critical for the observed self-organizing behavior of the protein that likely depend on specific hydrophobic interactions^42^.

While focused on a specific PrLD, these studies provide a framework for the in vitro characterization of condensate-forming biomolecules using complementary biophysical and structural techniques. The in vivo implications and prevalence of long-range molecule ordering of PrLD-containing proteins such as ELF3 are still to be fully explored. This may be a much more general phenomenon for other condensate-forming species. For example, the formation of discrete heterogeneous assemblies, or pleiomorphic ensembles, within the same droplet has been observed in vivo for scaffold proteins and signalling complexes, such as the PDGF receptor complex and the Wnt signalosome, allowing for the cooperative assembly of different components while retaining some conformational flexibility of the components^48,49^. Stacked structures in condensates have been observed for other IDR-containing proteins such as FLOE1, a plant water-sensory protein important for seed germination. Substitution of tyrosine and phenylalanine for serine residues in the aspartate-serine rich (DS) domain of FLOE1 resulted in gel formation of the mutant in vivo with ordered structures observed by TEM and 3-D tomography^50^. Phase separation and gelation of FLOE1 was driven by aromatic residues in the glutamine/proline/serine-rich (QPS) domain, in particular tyrosine residues.

Taken together, these experiments have allowed a detailed in vitro characterisation of ELF3 PrLD condensates using complementary biophysical and structural techniques that have not been previously applied to biological molecules in dilute and condensed phases. The molecular ordering observed in vitro for ELF3 PrLD may be a much more general phenomenon than previously appreciated and may help explain the variability in fluorescence recovery after photobleaching observed for many membraneless organelles^5,51,53^. An important and remaining challenge is to determine the physiological role of condensed species with different mobilities and whether long-range order and molecular stacking has a physiological role in ELF3 function. This will require further study including the re-engineering of the PrLD of ELF3 to alter LLPS and an examination of the effects in vivo at the cellular and phenotypic level.

## Experimental Methods

### Protein Expression and purification of ELF3PrLD and ELF3PrLD-GFP

ELF3 PrLD (Q7, residues 388-625, AT2G25930, *Arabidopsis thaliana* ecotype Columbia), Q0, Q20, ELF3 PrLD-GFP Q7, Q0 and Q20 were cloned into the expression vector pESPRIT2 using the Aat II and Not I sites as previously described^6^. To generate the GFP constructs the stop codon for ELF3 PrLD Q0, Q7 and Q20 was removed by site-directed mutagenesis using the QuikChange protocol (Agilent) and a GFP-tag was added to the C-terminus as previously described^6^. All proteins were overexpressed in *E. coli* BL21-CodonPlus-RIL cells (Agilent). Proteins used for phase diagrams, FRAP, AFM, SAXS, EM and X-ray diffraction experiments were expressed and purified as follows. Briefly, cells were grown in LB media supplemented with 50 μg ml^-1^ kanamycin and 35 μg ml^-1^ chloramphenicol at 37 °C and 120 rpm. At OD_600nm_ = 0.8, the temperature was lowered to 18 °C and protein expression was induced with 1 mM isopropyl-b-D-1-thiogalactopyranoside (IPTG). After 16 h, the cells were harvested by centrifugation at 5000 x g and 4 °C for 15 min. For all constructs, cells were resuspended in 100 mM Bis-Tris propane pH 9.4, 300 mM NaCl, 20 mM imidazole, 1 mM TCEP (tris(2-carboxyethyl)phosphine (TCEP) and EDTA-free protease inhibitors (ThermoFisher). Cells were lysed by sonication and the cell debris pelleted at 50000 x g and 4 °C for 30 min. The supernatant was applied to a 1 ml Ni-NTA column pre-equilibrated in resuspension buffer (lysis buffer without protease inhibitors), washed with resuspension buffer and high salt buffer (100 mM Bis-Tris propane pH 9.4, 1 M NaCl, 20 mM imidazole, 1 mM TCEP). The proteins were eluted in 100 mM Bis-Tris-propane pH 9.4, 300 mM NaCl, 300 mM imidazole and 1 mM TCEP. Protein purity was determined by SDS-PAGE, and the fractions of interest were pooled and dialysed for ~2 h at 4°C in 50 mM Bis-Tris-propane pH 9.4, 500 mM NaCl, 1 mM TCEP. Final protein concentration was 4-8 mg/ml for all samples. Bis-Tris-propane was used due to its wide buffering pH range. For the turbidity assays the Bis-Tris-propane was replaced with CAPS (N-cyclohexyl-3-aminopropanesulfonic acid) buffer, as the proteins were stable and soluble at this pH at protein concentrations. Purification was performed as for above, with the Bis-tris propane buffer replaced by CAPS pH 9.7. Final dialysis was performed against 50 mM CAPS pH 9.7, 200 mM NaCl, 1 mM TCEP and proteins diluted to ~0.4 mg/ml for turbidity assays.

### Transient expression in *Nicotiana benthamiana* for FRAP

The coding sequence of *ELF3Q0, ELF3Q7* and *ELF3Q20* were cloned into the GreenGate cloning system by digestion and ligation. For mVenus tagged constructs, GreenGate reactions with a β-estradiol inducible promoter were performed as previously described^53^. The *Agrobacterium tumefaciens* strain harbouring the pSOUP helper plasmid was transformed with the above-described plasmids. mVenus tagged fusion proteins were co-expressed with the p19 silencing repressor in epidermal leaf cells of *N. benthamiana.* Induction with β-estradiol was carried out as described ^54^.

### In vivo FRAP measurements and analysis of ELF3 condensates

FRAP measurements were carried out with a confocal laser scanning microscope (inverted LSM880, ZEISS) equipped with a 40x water objective (C-Apochromat, NA 1.2, ZEISS). mVenus was excited with an argon laser at 514 nm. For FRAP measurements, single condensates in or around the nucleus were chosen. Time series of 65 frames were acquired to observe fluorescence recovery. Acquisition times were approximately 300 ms per frame. Fluorescence recovery was analysed individually for each bleached condensate by drawing narrow ROIs ^55^. For normalisation, calculation of mobile fraction and half-life (τ½) a plugin written at the Stowers Institute for medical research (https://research.stowers.org/imagejplugins/index.html) was used in Fiji. 9-10 individual puncta were photobleached from different cells, and used in each analysis. FRAP curves were generated from all data and mean curves with S.D. are shown. Experiments were performed at room temperature.

### Light scattering turbidity assays

The light scattering assay was performed in a Cary 100 UV-vis spectrometer (Agilent Technologies UK Ltd., Stockport, UK) as previously reported^6^. Briefly, the absorbance at 440 nm was monitored for samples containing buffer alone (50 mM N-cyclohexyl-3-aminopropanesulfonic acid (CAPS) pH 9.7, 200 mM NaCl, 1 mM TCEP), ELF3 PrLD Q0, Q7 or Q20 (15 μM, ~ 0.4 mg/ml) in quartz cuvettes (path length 10 mm) with increasing temperature (10-40 °C; 1 °C min^-1^), and the spectra were normalized with respect to buffer alone. A transition temperature (Tm) was determined by fitting the spectrum with a 4-parameter sigmoidal equation using GraphPad Prism 9.4.0 and the sigmoidal 4PL equation, which is defined as y=A + (B-A)/(1+(Tm/x))^C^, where y is the normalised turbidity at 440 nm, x is the temperature in °C, A and B are bottom and top plateaus with the same units as y, C is the slope factor or Hill slope and Tm is the midpoint transition temperature. Reported values are from the curves shown in Fig. 2a with the filled blue squares for the first temperature increase from 10-40°C. Tm values were 31.2 ± 0.4°C (ELF3 PrLD Q0), 28.1 ± 0.3°C (ELF3 PrLD Q7) and 33.0 ± 1.5°C (ELF3 PrLD Q20). Due to the experimental set-up, where the highest temperature measured was 40 °C, the parameters extracted from the fitting, in particular for ELF3 PrLD Q20 should be considered as estimations as no plateau was reached and above 40 °C the proteins began to irreversibly precipitate. To monitor reversibility, the turbidity was monitored for an increasing temperature ramp (10 to 40 °C; 1 °C min^-1^) followed by decreasing the temperature (40 to 10 °C; 1 °C min^-1^) and this cycle was repeated twice in total for ELF3 PrLD Q0, Q7 and Q20.

### Phase diagrams

To generate phase diagrams for ELF3 PrLD Q0, Q7 and Q20 and ELF3 PrLD Q0-, Q7-, Q20 - GFP, the pH of the dialysis buffer (50 mM Bis-Tris propane pH 9.4, 500 mM NaCl and 1 mM TCEP) was gradually decreased by 0.2 pH units and at each pH the solution was allowed to equilibrate for at least 40 min. The solution was monitored for increased turbidity due to liquid droplet formation visually and by optical microscopic examination using the Olympus CKX41 bright field microscope with a Photometrics CoolSNAP cf2 camera at 20x magnification. All solutions were at 4 °C and kept in the cold room except for brief visualisation under a microscope at room temperature on cooled glass slides (4°C). For each phase diagram, measurements were performed at pH 9.5, 9.0, 8.8, 8.6, 8.4, 8.2, 8.0, 7.8, 7.6, 7.4 and 5 different protein concentrations (0.5, 1.0, 2.0, 4.0, 6.0 mg/ml).

### Temperature-induced LLPS images

Samples of ELF3 PrLD Q0, Q7 and Q20 at ~ 4 mg/ml in 50 mM Bis-Tris propane, pH 9.2, 500 mM NaCl and 1 mM TCEP (Q0 and Q7) or 50 mM Bis-Tris propane, pH 8.8, 500 mM NaCl and 1 mM TCEP (Q20), 500 mM NaCl, 1 mM TCEP were visualised at 4°C, 22°C and 4°C using an Olympus CKX41 bright field microscope with a Photometrics CoolSNAP cf2 camera at 20x magnification. 50 μl samples were cooled to ~ 4°C and 5 μl of solution was applied to a cooled glass slide and quickly imaged. The samples were then heated to 22°C in a heating block for ~5 min. to induce phase separation. 5μl of solution was applied to a room temperature glass slide and imaged. The tubes were then cooled to ~4°C for 15 min. and imaged as described.

### Confocal imaging and fluorescence microscopy in vitro

For droplet visualization and photobleaching experiments of ELF3 PrLD Q0-, Q7- and Q20-GFP proteins, liquid droplet formation was induced by dialysis as described above with GFP labelled ELF3 PrLD protein concentration of ~4 mg/ml and starting dialysis conditions of 50 mM Bis-Tris-propane pH 9.4, 500 mM NaCl with the pH gradually reduced in units of 0.2 until the solution became turbid (~pH 7.8). All dialysis steps were performed in a cold room at 4 °C. Once turbidity was observed, a 10 μl drop of solution was applied to a glass slide and all subsequent measurements performed at room temperature. The drop was covered with a cover slip and mounted and visualized using an objective-based total internal reflection fluorescence (TIRF) microscopy instrument composed of a Nikon Eclipse Ti, an azimuthal iLas2 TIRF illuminator (Roper Scientific), a 60x numerical aperture 1.49 TIRF objective lens followed by a 1.5x magnification lens and an Evolve 512 camera (Photometrics). For photobleaching experiments, droplets were allowed to adhere to the coverslip prior to photobleaching to minimize droplet movement during the experiment. Acquisition times were approximately 1 s per image for 30 s and then every 10 s for 270 s. Droplet size was ~2-5 μm with bleaching area of ~ 1 μm. Time-lapse images were acquired at 530 nm.

### Correlative AFM-fluorescence experiments

#### AFM-confocal fluorescence microscopy

AFM coupled to confocal fluorescence microscope was developed in-house, allowing AFM imaging and fluorescence recovery after photobleaching experiments on the same droplets (Supplementary Fig. S3a)^31^. AFM images were acquired using a Nanowizard 4 (JPK Instruments, Bruker) mounted on a Zeiss inverted optical microscope and equipped with a Tip Assisted Optics (TAO) module and a Vortis-SPM control unit. The AFM cantilever optical beam deflection system makes use of an infrared low-coherence light source (emission centred at 980 nm). A custom-made confocal microscope was coupled to the AFM using a supercontinuum laser (Rock-PP, Leukos) as laser source at 20 MHz equipped with an oil immersion objective with a 1.4 numerical aperture (Plan-Apochromat 100x, Zeiss). Fluorescence was collected after a pinhole of 100 *μ*m diameter size (P100D, Thorlabs) by an avalanche photodetector (SPCM-AQR-15, PerkinElmer) connected to an SPC-150 (Becker & Hickl) TCSPC card. An ET800sp short pass filter (Chroma) was used to filter out the light source of the AFM optical beam deflection system. The excitation laser power was measured after the objective at the sample level with a S170C microscope slide power sensor and a PM100 energy meter (both purchased from Thorlabs) and was set in all the experiments to 1 *μ*W. Confocal images and FRAP were acquired using a 488/10 nm excitation filter and a 525/39 nm emission filter. Simultaneous AFM/confocal images were collected with the AFM tip and confocal spot positions fixed and co-aligned while the sample was scanned using the TAO module. Full alignment was obtained using at first white field illumination. Then the fine-tuned was achieved measuring the increase of tip luminescence when aligned with the confocal spot. Finally, while imaging the sample, any mismatch between topography and confocal image was adjusted by correcting the tip position.

#### AFM-wide-field ffluorescence microscopy

AFM coupled wide-field fluorescence microscope was developed in-house, allowing AFM imaging and epifluorescence or Total Internal Reflection Fluorescence (TIRF) imaging (Supplementary Fig. S3b)^51,52^. and based on a Nanowizard 4 (JPK Instruments, Bruker) mounted on a Zeiss inverted optical microscope. The custom-made epifluorescence/TIRF microscope was coupled to the AFM using a LX 488-50 OBIS laser source (Coherent). We used an oil immersion objective with a 1.4 numerical aperture (Plan-Apochromat 100x, Zeiss). Fluorescence was collected with an EmCCD iXon Ultra897 (Andor) camera. The setup makes use of a 1.5x telescope to obtain a final imaging magnification of 150-fold, corresponding to a camera pixel size of 81 nm. An ET800sp short pass filter (Chroma) was used in the emission optical path to filter out the light source of the AFM optical beam deflection system. The excitation laser wavelength was centred at 488nm and the power was measured before the objective with a PM100 energy meter (purchased from Thorlabs) and was optimized in all the experiments in the range of 1-5 *μ*W. We used an acoustooptic tuneable filter (AOTFnc-400.650-TN, AA opto-electronics) to modulate the laser intensity and record fluorescence images using an ET525/50 nm (Chroma) as emission filter.

In both correlative AFM - TIRF / confocal, AFM images were acquired in QI mode with a scan size ranging from 10 *μ*m ×10 *μ*m to 50 *μ*m ×50 *μ*m and with 256×256 or 128/128 lines/pixels. Quantitative imaging mode (QI) was used to generate a force curve for each recorded pixel. Force curves were employed to evaluate droplets mechanical response. Typical force distance curves were recorded with a loading rate of ≈ 10 *μ*m /s, with large indentation cycle lengths ranging from 5 μm to 10 μm and maximal peak force of 1 – 1.5 nN. All parameters were optimized to provide image acquisition stability, considering sample low rigidity and having the cantilever immersed in a crowded environment with micrometric large droplets diffusing. Curves were then employed to extract all mechanical parameters for both liquid and gel phase constructs reported in Table 1, as well for the AFM images in Fig. 3 and Fig. 4. Gel phase constructs reported in Fig. 4, less dynamic on the coverslip and in solution, exhibited more stable force vs distance curves and could be imaged using a tip speed of 200 *μ*m/s, ensuring a faster acquisition time. Because of the micrometric height of the droplet comparable with the size of the AFM tip, only the curves acquired on the top of the droplets were considered, in order to avoid edge artifacts, non-negligible when the pyramidal tip facets are in contact with the droplets and present in the AFM images reported in Fig. 3 and Fig. 4c.

From the force curves on droplets in liquid phase, the stiffness keff was calculated based on a linear fit to the force vs indentation δ (Eq. 1)^32,33^.

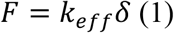

Indentation cycles performed on droplets in gel phase were treated using a Hertz contact model, leading to the evaluation of the Young’s modulus E (Eq. 2).

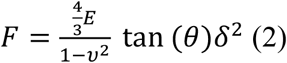

Where v is the Poisson’s ratio of 0.5, conventionally used for soft incompressible isotropic materials which are elastically deformed, and θ is the pyramidal tip angle. A comparison between indentation cycle performed on LLPS liquid phase (red) or gel condensates (blue) is shown in Fig. S4 where equations 1 and 2 have been used to fit the experimental data. The Young’s modulus images reported in Fig 4a exhibit a higher E compared to the values reported in table, which is likely due to the higher AFM tip speed (loading rate) employed for the acquisition of the force vs distance curves, suggesting a viscoelastic behaviour for all constructs in the gel phase.

Biolever mini (AC40TS, Olympus) and MSNL (Bruker) AFM cantilevers were purchased from Nano Bruker. Biolevers mini have a resonance frequency of ≈ 30 kHz in liquid, nominal stiffness of 0.1 N/m and a 7 *μ*m high tetrahedral tip. MSNL are V-shaped cantilevers with small tip radius (≈ 2 nm) suited for high resolution imaging, and nominal stiffness ranging from 0.01 to 0.6 N/m, depending on the cantilever. Images in Fig. 3 and Fig. 4 in the main manuscript were generated using AC40TS and MSNL AFM cantilevers, respectively. In all AFM experiments, the inverse optical lever sensitivity and lever stiffness of the cantilevers were calibrated using the “contact-free” method of the JPK AFM instruments, making use of a combination of a Sader and thermal methods^56,57^. For droplets in gel phase, we used the MSNL-E for Q0 and Q7 constructs and the MSNL-F for the Q20 ones, with nominal spring constant of 0.1 N/m and 0.6 N/m respectively, assuming *θ* = 22.5° to fit experimental data with equation (2).The fundamental resonance was used with a correction of 0.817 at room temperature and in liquid environment. Circular glass coverslips (25 mm diameter, 165 *μ*m thick) were purchased from Marienfeld. They were cleaned by a first cycle of sonication in 1 M KOH for 15 min, rinsed with deionized water 20 times, and finally subjected to a second sonication cycle in deionized water for 15 min. For measurements in the gel phase, GFP-labelled ELF3 PrLD Q0, Q7 and Q20 samples in buffer A (50 mM Bis-Tris propane, pH 9.4, 500 mM NaCl and 1 mM TCEP) at 4 mg/ml were dialysed stepwise against buffer B at pH 9.0 (100 mM Bis-Tris propane, 300 mM NaCl, 1 mM TCEP) for 45 minutes, followed by buffer B at pH 8.0 and finally buffer B at pH 7.6 for 15 minutes. The fast pH changes led to formation of spherical droplets with gel-like properties. The final protein concentration used for all measurements was ~2 mg/ml as calculated using A280 measurements on a NanoDrop (ThermoFisher) and adjusted by dilution with buffer B at pH 9.4. A 10 μL sample of protein was applied to a clean glass cover slip and mounted on the AFM. After incubation for 2 minutes to allow droplets to adhere to the surface, 300 μL of buffer A was added and finally imaged by correlative AFM-TIRF fluorescence (Fig. 4 in main manuscript). Total sample preparation time for AFM measurements was ~ 15-20 min. including sample slide preparation, cantilever tuning and image acquisition time (11 min. for a 256×256 pixel image). We have noted that droplets become more gel-like over time as they are allowed to adhere to the surface of the microscope slide.

From correlative confocal-AFM experiments, the LLPS liquid-like droplet contact angle can be evaluated. For a spherical droplet, it is given by^58^

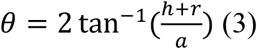

*θ* can be evaluated using the morphological information obtained from both confocal (parameter *a*) and AFM images (parameter *h*+*r*, maximal height of the droplet) as shown in Supplementary Fig. S5. Shadow vertically asymmetry in Fig. S5c can be partially due to the tetrahedral geometry of the probe as well as to the 10° cantilever inclination in respect to the glass coverslip (imposed by the AFM cantilever holder). Horizontal asymmetry can be partially due to the excitation beam not fully perpendicular to the glass coverslip and to a horizontal inclination of the cantilever. Contact angles for droplets were found in the range between 120° and 150° suggesting a low wettability associated to quasi-spherical droplet shapes as shown in Fig. 3 in main manuscript.

### Transmission electron microscopy experiments

Samples of gel phase Q0, Q7 and Q20 ELF3 PrLD were prepared by stepwise dialysis of proteins at 4 mg/ml in 50 mM Bis-Tris propane, pH 9.4, 500 mM NaCl and 1 mM TCEP. The pH was decreased in units of 0.2, with each dialysis step performed for 40 min at 4°C until pH 7.8. The sample became cloudy and then formed amorphous gel-like deposits, which were stored for at least 1hr at 4°C before image collection. The viscous gel was resuspended by fast pipetting to obtain a suspension of small gel pieces. Around 4 μl of sample was applied to the mica-carbon interface of a mica sheet with a layer of evaporated carbon film. The carbon film was floated off the mica sheet in ~600 μl of 2% (wt/vol) sodium silicotungstate. The sample was then transferred onto a 400-mesh copper TEM grid and air-dried. Images were taken on a Themo Fisher Technai F20 microscope operating at 200 kV. Images were acquired using a Thermo Fisher Ceta camera.

### SAXS measurements

SAXS experiments were performed at the European Synchrotron Radiation Facility (ESRF) on the BioSAXS beamline BM29^59,60^ for dilute phase, HPLC experiments and on beamline B21 at Diamond Light Source (SM26313-1, SM26314-2) for temperature ramp experiments^61^. For dilute phase experiments, an online HPLC system (Shimadzu) was attached directly to the sample inlet valve of the BM29 sample changer. Protein samples at 4 mg/ml (ELF3 PrLD Q0) or ~8 mg/ml (ELF3 PrLD Q7, Q20) were loaded into vials and automatically injected onto the column (Superose 6 3.2/300 Increase GE Healthcare) via an integrated syringe system. Buffers were degassed on-line and a flow rate of 0.05 ml/min at room temperature was used for all sample runs. For measurements of the dilute phase for ELF3 PrLD Q0, Q7 and Q20, 50 mM Bis-Tris propane pH 9.4, 500 mM NaCl and 1 mM TCEP was used for all samples. Prior to each run the column was equilibrated with at least 3 column volumes of buffer and the baseline monitored. All data from the run were collected using a sample to detector (Pilatus 2M Dectris) distance of 2.81 m corresponding to an q range of 0.008-0.45 Å^-1^. Due to column separation of the sample, some dilution effects will occur prior to measurement. 1800 frames (2 sec/frame) were collected. Initial data processing was performed automatically using the Dahu pipeline, generating radially integrated, calibrated and normalised 1-D profiles for each frame^62,63^. All frames were then further processed using Scatter IV^64^, briefly, the frames were dropped into the software and 50 – 100 frames selected for the background buffer. Using the heat plot a selection of similar frames (cyan in colour) were selected and merged. 30 frames corresponding to the highest protein concentration were merged and used for all further data processing and model fitting.

For the temperature ramp experiment for ELF3 PrLD Q7, samples were placed into the BioSAXS robot (Arinax, France) with both the sample changer and the vacuum cell both set at 4 °C. 50 μL of sample and matching buffer containing 50 mM CAPS, pH 9.7, 300 mM NaCl, 1mM TCEP were individually loaded and 10 frames of 1 second were taken. Temperature was raised in both the sample changer and the vacuum cell and incubated for 5 minutes before measurement. Temperature series of 4 °C, 12-15 °C, and 22-24 °C were taken. All frames were compared to the initial frame and matching frames were merged for buffer and samples. Scatter IV^10^ was then used to subtract buffer from sample and to generate Rg and plots.

### X-ray diffraction measurements

X-ray diffraction experiments for Q0, Q7 and Q20 ELF3PrLD were performed on beamline ID23-2 at the ESRF. Briefly, samples were purified as described above and dialysed directly against 50 mM BTP, pH 7.8, 250 mM NaCl and 1 mM TCEP at a concentyration of ~ 4 mg/ml for 3 h, after which time the protein formed an amorphous gel. The sample were spun down to collect the gel. The gel was directly mounted in a mesh litholoop (Molecular Dimensions). Diffraction was measured at room temperature. Diffuse powder rings were observed at ~4-5 Å, consistent with van der Waal’s distance between interacting molecules.

### Multi angle laser light scattering experiments

50 μl of ELF3 PrD Q0, Q7 or Q20 at a concentration of ~4-7mg/ml were loaded onto an S200 Increase size-exclusion column (Superdex 200 Increase10/300 GL, GE Healthcare) at a flow rate of 0.5 ml min-1. The column was pre-equilibrated with 50 mM Bis Tris propane at pH 9.4, 1 M NaCl, 1 mM TCEP and connected to a Hitachi Elite LaChrom UV detector and LAChrome Pump L-2130, a multi-angle laser light-scattering detector (DAWN HELEOS II, Wyatt Technology Corporation) and a refractive-index detector (Optilab T-rEX, Wyatt Technology Corporation). The data were processed with the ASTRA 6.1.7.17 software (Wyatt Technology Corporation).

## Supporting Information

Experimental materials and methods; supplementary figures for SAXS scattering curves and Kratky plots, additional X-ray protein sequences, additional phase diagrams for GFP labelled proteins. AFM and microscopy experimental set-up, linear and Hertz model fits for AFM data, additional AFM images, additional diffraction images, table of SAXS data collection statistics, additional references are provided as Supporting Information.

## Author Contributions

S.H., M.T., Y.S., V.I. S., A.D. and L.C. designed experiments, performed experiments and analysed data, J.K. and M.H.N. performed experiments and analysed data, W.L.L., A.P., A.M., A. G. and N.L. performed experiments, M.B. and P.-E. M. analysed data, P.W. designed experiments, C.Z. designed experiments and analysed data. The manuscript was written through contributions of all authors. All authors have given approval to the final version of the manuscript.

## Acknowledgment

We thank the B21 local contact Katsuaki Inoue from Diamond Light Source for his help during the SAXS temperature ramp experiments experiment. We thank Dr. Guy Schoehn for his support with EM experiments. We would like to thank the Partnership for Soft Condensed Matter (PSCM) at the ESRF for providing the lab space and equipment.

## Abbreviations

LLPS: liquid-liquid phase separation
PrLD: prion-like domain
IDR: intrinsically disordered region
ELF3: EARLY FLOWERING 3
SAXS: small angle X-ray scattering
AFM: atomic force microscopy
EM: electron microscopy

## References

1. Brangwynne, C.P., Eckmann, C.R., Courson, D.S., Rybarska, A., Hoege, C., Gharakhani, J., Jülicher, F., and Hyman, A.A. (2009). Germline P granules are liquid droplets that localize by controlled dissolution/condensation. Science 324, 1729–1732. 10.1126/science.1172046.

2. Feric, M., Vaidya, N., Harmon, T.S., Mitrea, D.M., Zhu, L., Richardson, T.M., Kriwacki, R.W., Pappu, R.V., and Brangwynne, C.P. (2016). Coexisting Liquid Phases Underlie Nucleolar Subcompartments. Cell 165, 1686–1697. 10.1016/j.cell.2016.04.047.

3. Riback, J.A., Zhu, L., Ferrolino, M.C., Tolbert, M., Mitrea, D.M., Sanders, D.W., Wei, M.-T., Kriwacki, R.W., and Brangwynne, C.P. (2020). Composition-dependent thermodynamics of intracellular phase separation. Nature 581, 209–214. 10.1038/s41586-020-2256-2.

4. Zhang, H., Elbaum-Garfinkle, S., Langdon, E.M., Taylor, N., Occhipinti, P., Bridges, A.A., Brangwynne, C.P., and Gladfelter, A.S. (2015). RNA Controls PolyQ Protein Phase Transitions. Mol Cell 60, 220–230. 10.1016/j.molcel.2015.09.017.

5. Shin, Y., Chang, Y.-C., Lee, D.S.W., Berry, J., Sanders, D.W., Ronceray, P., Wingreen, N.S., Haataja, M., and Brangwynne, C.P. (2018). Liquid Nuclear Condensates Mechanically Sense and Restructure the Genome. Cell 175, 1481–1491.e13. 10.1016/j.cell.2018.10.057.

6. Jh, J., Ad, B., S, H., Jr, K., M, G., D, D., Cs, S., X, L., E, P., F, G., et al. (2020). A prion-like domain in ELF3 functions as a thermosensor in Arabidopsis. Nature 585, 256–260. 10.1038/s41586-020-2644-7.

7. Franzmann, T.M., Jahnel, M., Pozniakovsky, A., Mahamid, J., Holehouse, A.S., Nüske, E., Richter, D., Baumeister, W., Grill, S.W., Pappu, R.V., et al. (2018). Phase separation of a yeast prion protein promotes cellular fitness. Science 359. 10.1126/science.aao5654.

8. Guillén-Boixet, J., Kopach, A., Holehouse, A.S., Wittmann, S., Jahnel, M., Schlüßler, R., Kim, K., Trussina, I.R.E.A., Wang, J., Mateju, D., et al. (2020). RNA-Induced Conformational Switching and Clustering of G3BP Drive Stress Granule Assembly by Condensation. Cell 181, 346–361.e17. 10.1016/j.cell.2020.03.049.

9. Iserman, C., Desroches Altamirano, C., Jegers, C., Friedrich, U., Zarin, T., Fritsch, A.W., Mittasch, M., Domingues, A., Hersemann, L., Jahnel, M., et al. (2020). Condensation of Ded1p Promotes a Translational Switch from Housekeeping to Stress Protein Production. Cell 181, 818–831.e19. 10.1016/j.cell.2020.04.009.

10. Alberti, S., Saha, S., Woodruff, J.B., Franzmann, T.M., Wang, J., and Hyman, A.A. (2018). A User’s Guide for Phase Separation Assays with Purified Proteins. J Mol Biol 430, 4806–4820. 10.1016/j.jmb.2018.06.038.

11. Protter, D.S.W., Rao, B.S., Treeck, B.V., Lin, Y., Mizoue, L., Rosen, M.K., and Parker, R. (2018). Intrinsically Disordered Regions Can Contribute Promiscuous Interactions to RNP Granule Assembly. Cell Reports 22, 1401–1412. 10.1016/j.celrep.2018.01.036.

12. Martin, E.W., and Holehouse, A.S. (2020). Intrinsically disordered protein regions and phase separation: sequence determinants of assembly or lack thereof. Emerging Topics in Life Sciences 4, 307–329. 10.1042/ETLS20190164.

13. Paloni, M., Bailly, R., Ciandrini, L., and Barducci, A. (2020). Unraveling Molecular Interactions in Liquid-Liquid Phase Separation of Disordered Proteins by Atomistic Simulations. J Phys Chem B 124, 9009–9016. 10.1021/acs.jpcb.0c06288.

14. Rauscher, S., and Pomès, R. (2017). The liquid structure of elastin. Elife 6. 10.7554/eLife.26526.

15. Murthy, A.C., Dignon, G.L., Kan, Y., Zerze, G.H., Parekh, S.H., Mittal, J., and Fawzi, N.L. (2019). Molecular interactions underlying liquid-liquid phase separation of the FUS low-complexity domain. Nat Struct Mol Biol 26, 637–648. 10.1038/s41594-019-0250-x.

16. Schuster, B.S., Dignon, G.L., Tang, W.S., Kelley, F.M., Ranganath, A.K., Jahnke, C.N., Simpkins, A.G., Regy, R.M., Hammer, D.A., Good, M.C., et al. (2020). Identifying sequence perturbations to an intrinsically disordered protein that determine its phase-separation behavior. Proc Natl Acad Sci U S A 117, 11421–11431. 10.1073/pnas.2000223117.

17. Dignon, G.L., Zheng, W., Kim, Y.C., Best, R.B., and Mittal, J. (2018). Sequence determinants of protein phase behavior from a coarse-grained model. PLoS Comput Biol 14, e1005941. 10.1371/journal.pcbi.1005941.

18. Das, S., Amin, A.N., Lin, Y.-H., and Chan, H.S. (2018). Coarse-grained residue-based models of disordered protein condensates: utility and limitations of simple charge pattern parameters. Phys Chem Chem Phys 20, 28558–28574. 10.1039/c8cp05095c.

19. Statt, A., Casademunt, H., Brangwynne, C.P., and Panagiotopoulos, A.Z. (2020). Model for disordered proteins with strongly sequence-dependent liquid phase behavior. J Chem Phys 152, 075101. 10.1063/1.5141095.

20. Dignon, G.L., Zheng, W., Kim, Y.C., and Mittal, J. (2019). Temperature-Controlled Liquid–Liquid Phase Separation of Disordered Proteins. ACS Cent. Sci. 5, 821–830. 10.1021/acscentsci.9b00102.

21. Elbaum-Garfinkle, S., Kim, Y., Szczepaniak, K., Chen, C.C.-H., Eckmann, C.R., Myong, S., and Brangwynne, C.P. (2015). The disordered P granule protein LAF-1 drives phase separation into droplets with tunable viscosity and dynamics. Proc Natl Acad Sci U S A 112, 7189–7194. 10.1073/pnas.1504822112.

22. Burke, K.A., Janke, A.M., Rhine, C.L., and Fawzi, N.L. (2015). Residue-by-Residue View of In Vitro FUS Granules that Bind the C-Terminal Domain of RNA Polymerase II. Mol Cell 60, 231–241. 10.1016/j.molcel.2015.09.006.

23. Kroschwald, S., Munder, M.C., Maharana, S., Franzmann, T.M., Richter, D., Ruer, M., Hyman, A.A., and Alberti, S. (2018). Different Material States of Pub1 Condensates Define Distinct Modes of Stress Adaptation and Recovery. Cell Rep 23, 3327–3339. 10.1016/j.celrep.2018.05.041.

24. Riback, J.A., Katanski, C.D., Kear-Scott, J.L., Pilipenko, E.V., Rojek, A.E., Sosnick, T.R., and Drummond, D.A. (2017). Stress-Triggered Phase Separation Is an Adaptive, Evolutionarily Tuned Response. Cell 168, 1028–1040.e19. 10.1016/j.cell.2017.02.027.

25. Tajima, T., Oda, A.O., Nakagawa, M., Kamada, H., and Mizoguchi, T. (2007). Natural variation of polyglutamine repeats of a circadian clock gene ELF 3 in Arabidopsis. In.

26. Undurraga, S.F., Press, M.O., Legendre, M., Bujdoso, N., Bale, J., Wang, H., Davis, S.J., Verstrepen, K.J., and Queitsch, C. (2012). Background-dependent effects of polyglutamine variation in the Arabidopsis thaliana gene ELF3. Proc Natl Acad Sci U S A 109, 19363–19367. 10.1073/pnas.1211021109.

27. McSwiggen, D.T., Mir, M., Darzacq, X., and Tjian, R. (2019). Evaluating phase separation in live cells: diagnosis, caveats, and functional consequences. Genes Dev 33, 1619–1634. 10.1101/gad.331520.119.

28. Park, S., Barnes, R., Lin, Y., Jeon, B., Najafi, S., Delaney, K.T., Fredrickson, G.H., Shea, J.-E., Hwang, D.S., and Han, S. (2020). Dehydration entropy drives liquid-liquid phase separation by molecular crowding. Commun Chem 3, 1–12. 10.1038/s42004-020-0328-8.

29. Lin, Y., McCarty, J., Rauch, J.N., Delaney, K.T., Kosik, K.S., Fredrickson, G.H., Shea, J.-E., and Han, S. Narrow equilibrium window for complex coacervation of tau and RNA under cellular conditions. eLife 8, e42571. 10.7554/eLife.42571.

30. Quiroz, F.G., and Chilkoti, A. (2015). Sequence heuristics to encode phase behaviour in intrinsically disordered protein polymers. Nat Mater 14, 1164–1171. 10.1038/nmat4418.

31. Fernandes, T.F.D., Saavedra-Villanueva, O., Margeat, E., Milhiet, P.-E., and Costa, L. (2020). Synchronous, Crosstalk-free Correlative AFM and Confocal Microscopies/Spectroscopies. Scientific Reports 10, 7098. 10.1038/s41598-020-62529-3.

32. Costa, L., Li-Destri, G., Pontoni, D., Konovalov, O., and Thomson, N.H. (2017). Liquid–Liquid Interfacial Imaging Using Atomic Force Microscopy. Advanced Materials Interfaces 4, 1700203. 10.1002/admi.201700203.

33. Chan, D.Y.C., Dagastine, R.R., and White, L.R. (2001). Forces between a Rigid Probe Particle and a Liquid Interface. J Colloid Interface Sci 236, 141–154. 10.1006/jcis.2000.7400.

34. Martin, E.W., Hopkins, J.B., and Mittag, T. (2021). Chapter Seven - Small-angle X-ray scattering experiments of monodisperse intrinsically disordered protein samples close to the solubility limit. In Methods in Enzymology Liquid-Liquid Phase Coexistence and Membraneless Organelles., C. D. Keating, ed. (Academic Press), pp. 185–222. 10.1016/bs.mie.2020.07.002.

35. Martin, E.W., Harmon, T.S., Hopkins, J.B., Chakravarthy, S., Incicco, J.J., Schuck, P., Soranno, A., and Mittag, T. (2021). A multi-step nucleation process determines the kinetics of prion-like domain phase separation. Nat Commun 12, 4513. 10.1038/s41467-021-24727-z.

36. Shirahama, T., and Cohen, A.S. (1965). Structure of Amyloid Fibrils after Negative Staining and High-resolution Electron Microscopy. Nature 206, 737–738. 10.1038/206737a0.

37. Meister, A., and Blume, A. (2017). (Cryo)Transmission Electron Microscopy of Phospholipid Model Membranes Interacting with Amphiphilic and Polyphilic Molecules. Polymers (Basel) 9, 521. 10.3390/polym9100521.

38. Hope, M.J., Bally, M.B., Webb, G., and Cullis, P.R. (1985). Production of large unilamellar vesicles by a rapid extrusion procedure: characterization of size distribution, trapped volume and ability to maintain a membrane potential. Biochim Biophys Acta 812, 55–65. 10.1016/0005-2736(85)90521-8.

39. Hughes, M.P., Sawaya, M.R., Boyer, D.R., Goldschmidt, L., Rodriguez, J.A., Cascio, D., Chong, L., Gonen, T., and Eisenberg, D.S. (2018). Atomic structures of low-complexity protein segments reveal kinked β sheets that assemble networks. Science 359, 698–701. 10.1126/science.aan6398.

40. Bellingham, C.M., Lillie, M.A., Gosline, J.M., Wright, G.M., Starcher, B.C., Bailey, A.J., Woodhouse, K.A., and Keeley, F.W. (2003). Recombinant human elastin polypeptides self-assemble into biomaterials with elastin-like properties. Biopolymers 70, 445–455. 10.1002/bip.10512.

41. Lyons, R.E., Nairn, K.M., Huson, M.G., Kim, M., Dumsday, G., and Elvin, C.M. (2009). Comparisons of recombinant resilin-like proteins: repetitive domains are sufficient to confer resilin-like properties. Biomacromolecules 10, 3009–3014. 10.1021/bm900601h.

42. Garcia Garcia, C., and Kiick, K.L. (2019). Methods for producing microstructured hydrogels for targeted applications in biology. Acta Biomater 84, 34–48. 10.1016/j.actbio.2018.11.028.

43. Schuster, B.S., Reed, E.H., Parthasarathy, R., Jahnke, C.N., Caldwell, R.M., Bermudez, J.G., Ramage, H., Good, M.C., and Hammer, D.A. (2018). Controllable protein phase separation and modular recruitment to form responsive membraneless organelles. Nat Commun 9, 2985. 10.1038/s41467-018-05403-1.

44. Shin, Y., Berry, J., Pannucci, N., Haataja, M.P., Toettcher, J.E., and Brangwynne, C.P. (2017). Spatiotemporal Control of Intracellular Phase Transitions Using Light-Activated optoDroplets. Cell 168, 159–171.e14. 10.1016/j.cell.2016.11.054.

45. Tompa, P., and Fuxreiter, M. (2008). Fuzzy complexes: polymorphism and structural disorder in protein-protein interactions. Trends Biochem Sci 33, 2–8. 10.1016/j.tibs.2007.10.003.

46. Wu, H., and Fuxreiter, M. (2016). A Structure and Dynamics Continuum of Higher-Order Assemblies: Amyloids, Signalosomes and Granules. Cell 165, 1055–1066. 10.1016/j.cell.2016.05.004.

47. Hughes, M.P., Goldschmidt, L., and Eisenberg, D.S. (2021). Prevalence and species distribution of the low-complexity, amyloid-like, reversible, kinked segment structural motif in amyloid-like fibrils. Journal of Biological Chemistry 297, 101194. 10.1016/j.jbc.2021.101194.

48. Korkmazhan, E., Tompa, P., and Dunn, A.R. (2021). The role of ordered cooperative assembly in biomolecular condensates. Nat Rev Mol Cell Biol 22, 647–648. 10.1038/s41580-021-00408-z.

49. Mayer, B.J., Blinov, M.L., and Loew, L.M. (2009). Molecular machines or pleiomorphic ensembles: signaling complexes revisited. Journal of Biology 8, 81. 10.1186/jbiol185.

50. Dorone, Y., Boeynaems, S., Flores, E., Jin, B., Hateley, S., Bossi, F., Lazarus, E., Pennington, J.G., Michiels, E., De Decker, M., et al. (2021). A prion-like protein regulator of seed germination undergoes hydration-dependent phase separation. Cell 184, 4284–4298.e27. 10.1016/j.cell.2021.06.009.

51. Dahmane, S., Doucet, C., Le Gall, A., Chamontin, C., Dosset, P., Murcy, F., Fernandez, L., Salas, D., Rubinstein, E., Mougel, M., et al. (2019). Nanoscale organization of tetraspanins during HIV-1 budding by correlative dSTORM/AFM. Nanoscale 11, 6036–6044. 10.1039/c8nr07269h.

52. Vial, A., Taveneau, C., Costa, L., Chauvin, B., Nasrallah, H., Godefroy, C., Dosset, P., Isambert, H., Ngo, K.X., Mangenot, S., et al. (2021). Correlative AFM and fluorescence imaging demonstrate nanoscale membrane remodeling and ring-like and tubular structure formation by septins. Nanoscale 13, 12484–12493. 10.1039/d1nr01978c.

53. Lampropoulos, A., Sutikovic, Z., Wenzl, C., Maegele, I., Lohmann, J.U., and Forner, J. (2013). GreenGate - A Novel, Versatile, and Efficient Cloning System for Plant Transgenesis. PLOS ONE 8, e83043. 10.1371/journal.pone.0083043.

54. Bleckmann, A., Weidtkamp-Peters, S., Seidel, C.A.M., and Simon, R. (2010). Stem Cell Signaling in Arabidopsis Requires CRN to Localize CLV2 to the Plasma Membrane. Plant Physiology 152, 166–176. 10.1104/pp.109.149930.

55. Schindelin, J., Arganda-Carreras, I., Frise, E., Kaynig, V., Longair, M., Pietzsch, T., Preibisch, S., Rueden, C., Saalfeld, S., Schmid, B., et al. (2012). Fiji: an open-source platform for biological-image analysis. Nat Methods 9, 676–682. 10.1038/nmeth.2019.

56. Sader, J.E., Borgani, R., Gibson, C.T., Haviland, D.B., Higgins, M.J., Kilpatrick, J.I., Lu, J., Mulvaney, P., Shearer, C.J., Slattery, A.D., et al. (2016). A virtual instrument to standardise the calibration of atomic force microscope cantilevers. Review of Scientific Instruments 87, 093711. 10.1063/1.4962866.

57. Proksch, R., Schäffer, T.E., Cleveland, J.P., Callahan, R.C., and Viani, M.B. (2004). Finite optical spot size and position corrections in thermal spring constant calibration. Nanotechnology 15, 1344–1350. 10.1088/0957-4484/15/9/039.

58. Nadargi, D., Latthe, S., Hirashima, H., and Rao, A. (2009). Studies on Rheological Properties of Methyltriethoxysilane (MTES) Based Flexible Superhydrophobic Silica Aerogels. Microporous and Mesoporous Materials 117, 617–626. 10.1016/j.micromeso.2008.08.025.

59. Pernot, P., Round, A., Barrett, R., De Maria Antolinos, A., Gobbo, A., Gordon, E., Huet, J., Kieffer, J., Lentini, M., Mattenet, M., et al. (2013). Upgraded ESRF BM29 beamline for SAXS on macromolecules in solution. Journal of synchrotron radiation 20, 660–664. 10.1107/S0909049513010431.

60. Tully, M.D., Kieffer, J., Brennich, M.E., Cohen Aberdam, R., Florial, J.B., Hutin, S., Oscarsson, M., Beteva, A., Popov, A., Moussaoui, D., et al. (2023). BioSAXS at European Synchrotron Radiation Facility – Extremely Brilliant Source: BM29 with an upgraded source, detector, robot, sample environment, data collection and analysis software. J Synchrotron Rad 30, 258–266. 10.1107/S1600577522011286.

61. Cowieson, N.P., Edwards-Gayle, C.J.C., Inoue, K., Khunti, N.S., Doutch, J., Williams, E., Daniels, S., Preece, G., Krumpa, N.A., Sutter, J.P., et al. (2020). Beamline B21: high-throughput small-angle X-ray scattering at Diamond Light Source. J Synchrotron Rad 27, 1438–1446. 10.1107/S1600577520009960.

62. Incardona, M.F., Bourenkov, G.P., Levik, K., Pieritz, R.A., Popov, A.N., and Svensson, O. (2009). EDNA: a framework for plugin-based applications applied to X-ray experiment online data analysis. J Synchrotron Radiat 16, 872–879. 10.1107/S0909049509036681.

63. Kieffer, J., Brennich, M., Florial, J.B., Oscarsson, M., De Maria Antolinos, A., Tully, M., and Pernot, P. (2022). New data analysis for BioSAXS at the ESRF. J Synchrotron Radiat 29, 1318–1328. 10.1107/S1600577522007238.

64. Tully, M.D., Tarbouriech, N., Rambo, R.P., and Hutin, S. (2021). Analysis of SEC-SAXS data via EFA deconvolution and Scatter. JoVE (Journal of Visualized Experiments), e61578. 10.3791/61578.

