## Supplemental Information for "Phase separation and molecular ordering of the prion-like domain of the thermosensory protein EARLY FLOWERING 3"

### Supplementary Figures

**Figure S1.** Sequence and phase diagrams for ELF3 PrLD GFP constructs. **A)** *Arabidopsis thaliana* EARLY FLOWERING 3 PrLD constructs with varying polyQ regions used in the in vitro experiments. The 6x histidine tag and TEV protease cleavage site is shown in blue, the ELF3 PrLD sequence is in black with the polyQ expansion region shown as a star. The polyQ PrLD constructs used in this study correspond to Q0 (no glutamines), Q7 and Q20 as shown schematically as a triangle above the sequence. The GFP tag used in the fluorescence microscopy experiments is shown in green. **B)** Phase diagram of GFP labelled ELF3 PrLD constructs. Clear empty circles are dilute phase, blue filled circles liquid droplets and grey filled circles are precipitate/gel.

A

MGHHHHHHDYDIPTTENLYFQGRRLLPSEFLVKPPLPHVVVKQRGDSEKTDQHKMESSAENVVGRLSNQG

HHQQSNYMPFANNPPASPAPNGYCFPPQPPPSGNHQQWLIPVMSPSEGLIYKPHPGMAHTGHYGGYYGHY

QQQQQQQQQQQQQQQQQQQQ 20

QQQQQQQ 7

MPTPMVMPQYHPGMGFPPPGNGYFPPYGMMPTIMNPYCSS*PNEQMNQFGHPGNLQNTQQQQQRSDNEPAP

QQQQQPTKSYPRARKSRQGSTGSSPSGPQGISGSKSFRPFAAVDE-GFP

B

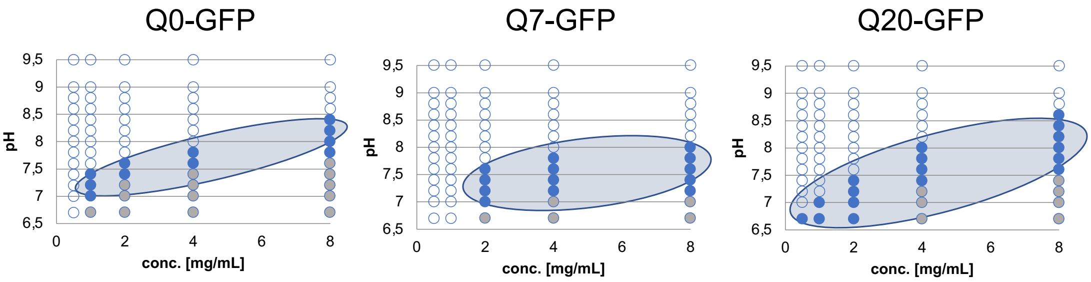

**Figure S2.** Experimental set-up and analytical consideration for AFM experiments. **A)** AFM/confocal (Figure 3) and **B)** AFM/TIRF (Figure 4) experimental set-up. **C**) Example of force versus distance measurements for ELF3-PrLD Q20 with curve fits using a linear fit (liquid samples, red curve) and Hertz fit for calculation of the Young's modulus (gel samples, blue curve). Data given in Table 1**. D-F)** Example of contact angle calculations (Eq. 3) using correlative confocal-AFM demonstrating low wettability of gel phase droplets. **1)** droplet geometry onto the glass substrate. **2)** and **3)** AFM topography and confocal images, respectively, used to evaluate the contact angle of ≈ 140°: the maximal height of the droplet *h+r* is obtained from the AFM image (**2**), whereas *a* is obtained from the confocal image (**3**). Therefore, the AFM image provides the droplet height, whereas the confocal image provides the diameter of the circular droplet surface in contact with the glass substrate. The shadow present in the confocal image is due to a lower fluorescence of the emitters that stick to the AFM tip. This shadow corresponds with the droplet edges and is due to the side of the pyramidal AFM tip in contact with the droplet, resulting in a vertical position of the AFM tip which is higher than the substrate position, which is the focal plane of the confocal microscope. The fluorescence background due to the emitters coating the AFM tip vanishes in correspondence with a height variation during tip scanning because the AFM tip gets far from the focal plane.

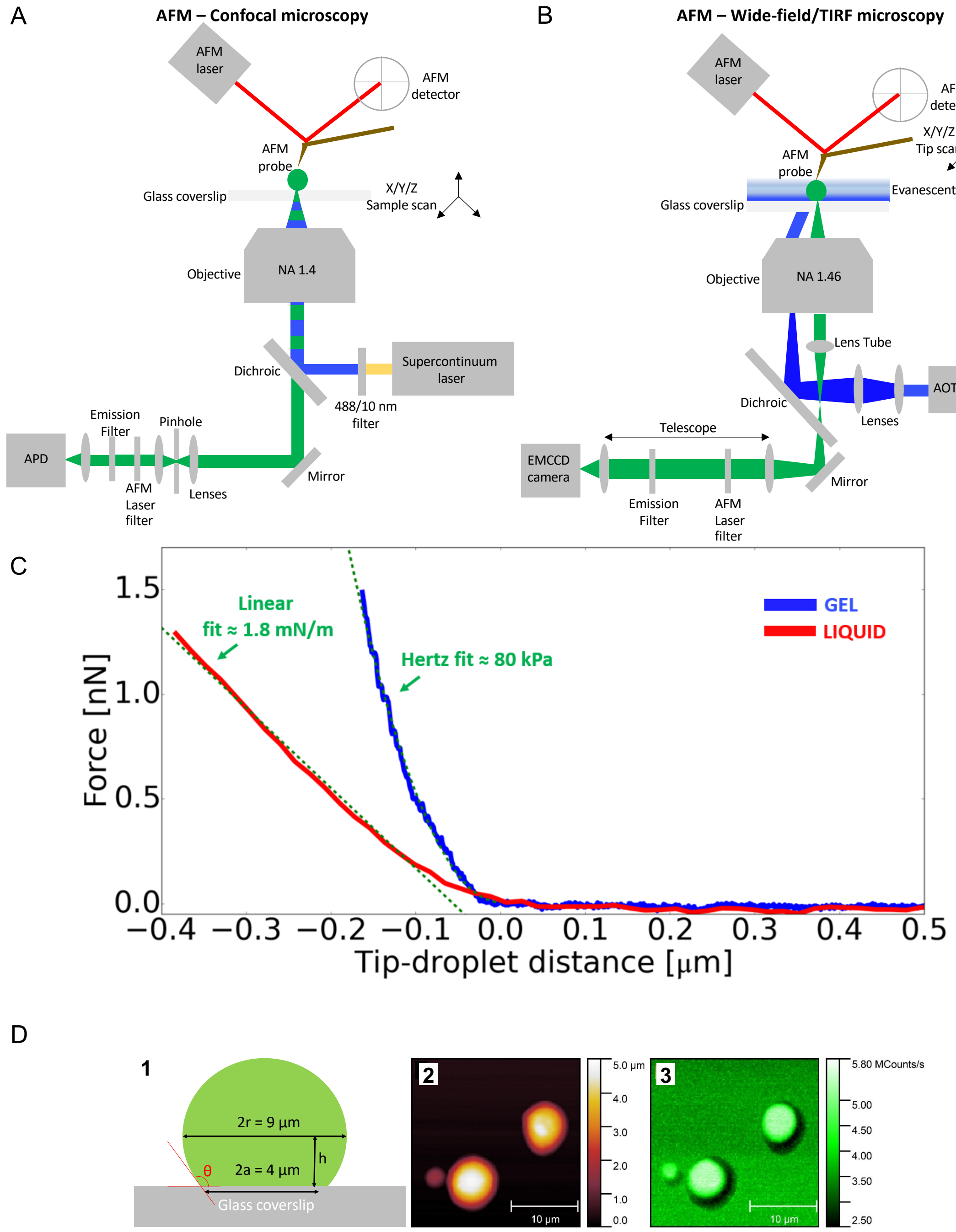

**Figure S3.** AFM images of ELF3-GFP PrLD samples in gel phase. As for the images reported in Fig. 4, here constructs exhibit a stacked or layered structure with step-like height profiles ranging from ~20 nm to 200 nm. **A**) Q0 construct, scale bar is 1 micron and height are shown as gray scale as indicated. **B**) Q7 labelled as per (**A**). **C**) Q20 construct labelled as per (**A**). The layered structure was not observed in the droplets in liquid phase (Fig. 3).

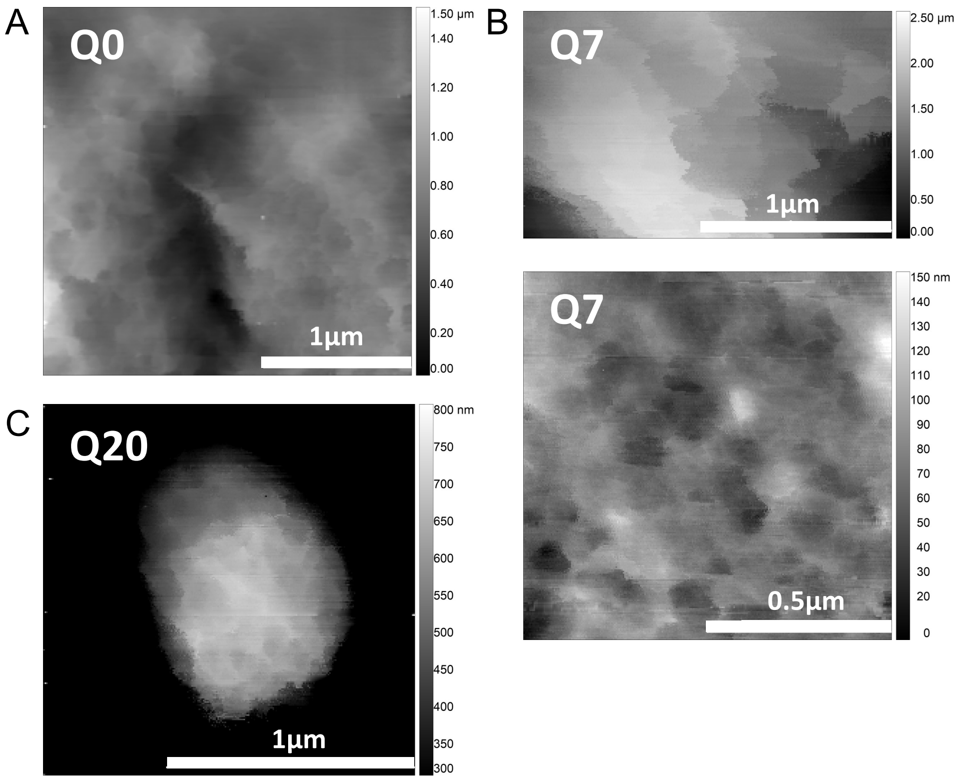

**Figure S4.** Size exclusion multiangle laser light scattering (SEC-MALLS) for Q0, Q7 and Q20 constructs. All peaks eluted after the void volume of the S200 column (GE Healthcare) and exhibited a monomodal particle-size distribution with molar masses of ~ 1000 kDa, corresponding to a large 25-30-mer species in the dilute phase. UV traces are in blue and molecular weight are show in yellow.

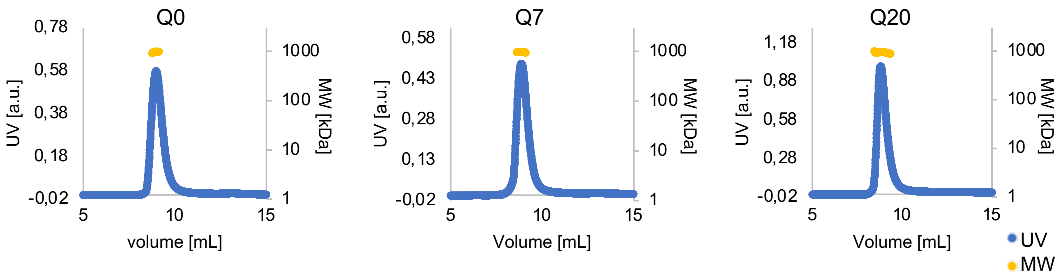

976.2 kDa    967.6 kDa    936.2 kDa

**Figure S5.** SAXS scattering curves with increasing temperature. ELF3 PrLD in the condensed phase **A)** Q0 (green) at temperatures 4, 12 and 24 °C, **B)** Q7 (pink) at temperatures 4, 15 and 22 °C and **C)** Q20 (blue) at temperatures 4, 12 and 24 °C. The left plots show the intensity as a function of q, where q=(4π/λ sin θ) and the right plots show the corresponding total intensity plot (I*q vs q) which emphasizes the structure factor peak. For both plots the data has been trimmed to q = 0.06 Å to focus on the area of interest.

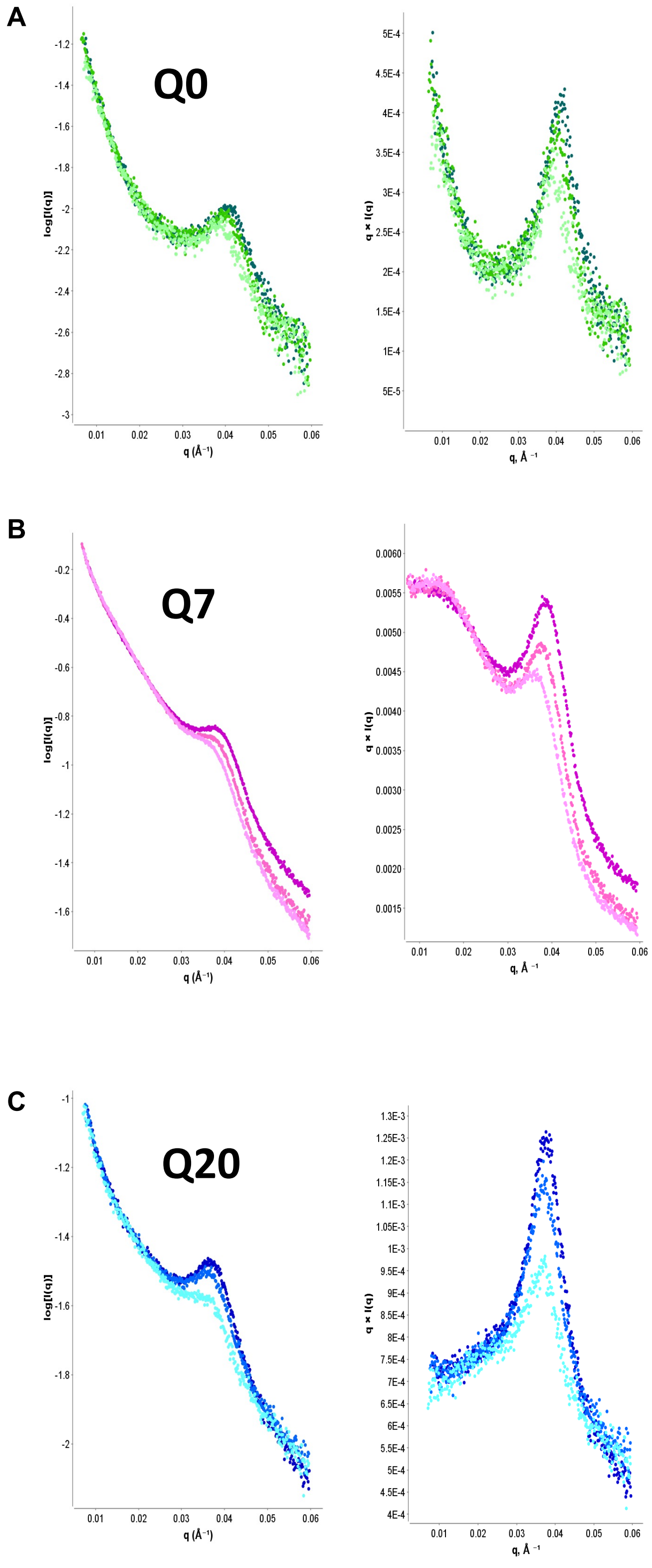

**Figure S6.** X-ray scattering for ELF3-PrLD hydrogels. Empty loop scattering in shown at top left. Q0 and Q7 show only diffuse solvent scattering whereas Q20 exhibits a powder diffraction ring indicated by the blue arrow. All images taken at room temperature; resolution rings are indicated.

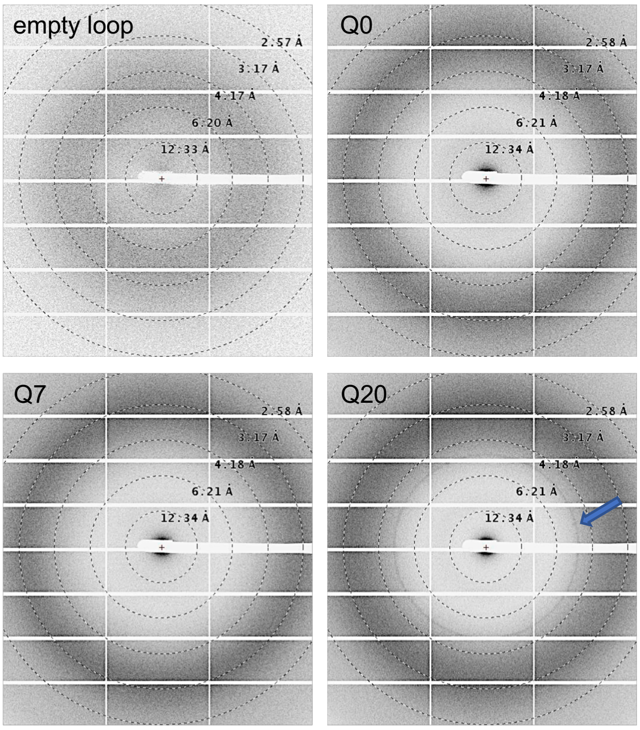

**Table S1. Summary of SAXS data collections. Data collected at Diamond Light Source (DLS) and the European Synchrotron Radiation Facility (ESRF).**

| **Data Collection Parameters** | DLS | ESRF |
|---|---|---|
| q-range ($Å^{-1}$) | 0.0026-0.34 | 0.007-0.5 |
| Sample-to-detector distance (m) | 3.7 | 2.6 |
| Concentration range (mg/mL) | 1-6 | 1-6 |
| Temperature (K) | 277-300 | 277-300 |
| Detector | Eiger 4M (Dectris) | Pilatus 1M/Pilatus P3-2M |
| Flux (photons/s) | $2*10^{12}$ | $1.4*10^{12}/1*10^{13}$ |
| Beam size (μm) | 1102*240 | 700*700 /500*200 |

| **Structural parameters (heat ramp)** | **Q0 (4 °C)** | **Q0 (12 °C)** | **Q0 (24 °C)** |
|---|---|---|---|
| I0 (kDa) [from Guinier] | 0.06 | 0.021 | 0.013 |
| Rg (Å) [from Guinier] | 129.1 | 67.5 | 44.1 |
| $q_{min}Rg$ - $q_{max}Rg$ used for Guinier | $5*10^{-4}-1*10^{-4}$ | $2.4*10^{-4}-3.7*10^{-4}$ | $4*10^{-5}-9*10^{-5}$ |
| q-range peak value($Å^{-1}$) | 0.039 | 0.0395 | 0.0405 |
| domain spacing, d (Å) (d=2π/q) | 161 | 159 | 155 |

| **Structural parameters (heat ramp)** | **Q7 (4 °C)** | **Q7 (15 °C)** | **Q7 (22 °C)** |
|---|---|---|---|
| I0 (kDa) [from Guinier] | 0.74 | 0.61 | 0.46 |
| Rg (Å) [from Guinier] | 97 | 94 | 97.6 |
| $q_{min}Rg$ - $q_{max}Rg$ used for Guinier | $1.4*10^{-4}-1.8*10^{-4}$ | $1.35*10^{-4}-1.9*10^{-4}$ | $1.3*10^{-4}-1.8*10^{-4}$ |
| q-range peak value($Å^{-1}$) | 0.0385 | 0.0375 | 0.0385 |
| domain spacing, d (Å) | 163 | 167 | 163 |

| **Structural parameters (heat ramp)** | **Q20 (4 °C)** | **Q20 (12 °C)** | **Q20 (24 °C)** |
|---|---|---|---|
| I0 (kDa) [from Guinier] | 0.075 | 0.087 | 0.1 |
| Rg (Å) [from Guinier] | 94.6 | 104.7 | 105.1 |
| $q_{min}Rg$ - $q_{max}Rg$ used for Guinier | $1.1*10^{-4}-1.9*10^{-4}$ | $1*10^{-4}-1.6*10^{-4}$ | $9*10^{-5}-1.6*10^{-4}$ |
| q-range peak value($Å^{-1}$) | 0.037 | 0.037 | 0.0375 |
| domain spacing, d (Å) | 170 | 170 | 167 |

| **Structural parameters (HPLC) (dilute phase)** | **Q0** | **Q7** | **Q20** |
|---|---|---|---|
| I0 (kDa) [from Guinier] | 213 | 369 | 274 |
| Rg (Å) [from Guinier] | 72.4 | 73.4 | 75.6 |
| $q_{min}Rg$ - $q_{max}Rg$ used for Guinier | $1.1*10^{-2}-1.8*10^{-2}$ | $1.3*10^{-2}-1.8*10^{-2}$ | $1.3*10^{-2}-1.7*10^{-2}$ |
| Volume (Å3) | 1.8*106 | 1.9*106 | 2.0*106 |
| $D_{max}$ (Å) | 272 | 273 | 294 |
| q-range peak value($Å^{-1}$) | N/A | N/A | N/A |
| Calculated MW | 750 | 820 | 1100 |
| Calculated theoretical MW monomer (kDa): | 28.26 | 29.15 | 30.82 |

| **Software employed** | | |
|---|---|---|
| Primary data reduction: | B21 autoprocessing pipeline | BM29 autoprocessing pipeline |
| Data processing | Scatter IV | |
